# Biallelic Loss of Molecular Chaperone Molecule AIP Results in a Novel Severe Multisystem Disease Defined by Defective Proteostasis

**DOI:** 10.1101/2024.08.08.604602

**Authors:** Márta Korbonits, Xian Wang, Sayka Barry, Chung Thong Lim, Oniz Suleyman, Stefano De Tito, Nazia Uddin, Maria Lillina Vignola, Charlotte Hall, Laura Perna, J. Paul Chapple, Gabor Czbik, Sian M Henson, Valle Morales, Katiuscia Bianchi, Viðar Örn Eðvarðsson, Kristján Ari Ragnarsson, Viktoría Eir Kristinsdóttir, Anne Debeer, Yoeri Sleyp, Rena Zinchenko, Glenn Anderson, Michael Duchen, Kritarth Singh, Chih Yao Chung, Yu Yuan, Sandip Patel, Ezra Aksoy, Artem O. Borovikov, Hans Tómas Björnsson, Hilde Van Esch, Sharon Tooze, Caroline Brennan, Oliver Haworth

## Abstract

Children born with deleterious biallelic variants of the chaperone aryl hydrocarbon receptor interacting protein (AIP) have a novel pediatric metabolic disease presenting a severe, complex clinical phenotype characterized by failure to develop following birth. Analysis of *Aip* knockout mouse embryonic fibroblasts and patient-derived dermal fibroblasts revealed that AIP was required to support proteostasis; including proteasome activity, induction of autophagy and lysosome function. aip knockout zebrafish, recapitulated the phenotype of the children; dying at an early stage of development when autophagy is required to adapt to periods of starvation. Our results demonstrate that AIP plays a crucial role in initiating autophagy and maintaining proteostasis *in vitro* and *in vivo*.

**One Sentence Summary:** Homozygous loss of the chaperone AIP results in a novel pediatric disease exhibiting multiple features of a lysosomal storage disease.

## Introduction

The proteostasis network coordinates protein synthesis, folding, disaggregation and degradation (Hartl, Bracher and Hayer-Hartl, 2011). The Ubiquitin-Proteasome System (UPS) and autophagy pathways mediate protein degradation, with molecular chaperones triaging substrates to them (Joshi *et al*., 2018). Disruption of proteostasis is associated with cancer, neurodegenerative, metabolic and many other diseases (Wolff, Weissman and Dillin, 2014). Autophagy is an evolutionary conserved catabolic process that is integral for maintaining proteostasis and wider cellular homeostasis in response to the dynamic metabolic demands of cells and organisms and essential for organisms adapting to periods of fasting (Levine and Kroemer, 2019; Efeyan *et al*., 2013; Kuma *et al*., 2004; Komatsu *et al*., 2005; Maweed Suzan Attia, 2021). Crucial to autophagic recycling of proteins are lysosomes which are at the center of nutrient catabolism within the cell, recycling nutrients from within the cell and enabling mTOR to detect and respond to nutrient availability within the cell (Goul, Peruzzo and Zoncu, 2023).

Aryl hydrocarbon receptor interacting protein (AIP) is a highly conserved ubiquitously expressed molecular chaperone (Trivellin and Korbonits, 2011). Initially described as a co-chaperone to heat shock proteins HSP90 and HSC70 and shown to support the deubiquitinase UCHL1 (Sun *et al*., 2019). Heterozygote loss-of-function germline variants in *AIP* predispose to the development of pituitary adenomas (Loughrey and Korbonits, 2019; Vierimaa *et al*., 2006). However, the metabolic processes that AIP supports are still unknown.

Based upon previous experimental evidence, it was predicted that deleterious biallelic variants of AIP would be incompatible with life, as loss of AIP resulted in embryonic lethality in mice, *Drosophila* and *C. elegans* (Lin *et al*., 2007; Aflorei *et al*., 2018; Chen *et al*., 2017). We now describe patients born with biallelic *AIP* variants with a novel, severe, complex pediatric disease that shares similarities with lysosomal storage diseases (LSDs), providing evidence that AIP is fundamental in maintaining proteostasis by supporting autophagy, lysosomes and the proteasome using *Aip* knockout (KO) mouse embryonic fibroblasts (*Aip* KO MEFs), homozygous *AIP* mutant patient derived dermal fibroblasts (AIPd-PDFs) and an *in vivo* zebrafish model of AIP deficiency.

## Results

We have identified five patients form three different countries with a uniform and unique phenotype representing a novel disease: hyperthermia (non-infectious/non-inflammatory), tachycardia, hypercalcemia, severe diarrhea and failure-to-thrive (**Figure 1A**, **Figure S1A**), despite being given state-of-the-art care and feeding support in pediatric intensive care units (**Figure 1B**). The initial two identified patients were first cousins from a consanguineous family (Family A) and died with low body weight and heart failure under the age of 12 months. Whole exome sequencing was performed and a biallelic variant was identified in both patients (Family A Member 1 and 2 (FAM1, FAM2)) in the *AIP* gene: c.62G>A; p.Gly21Asp (NM_003977.4) (**Figure 1A**) (**Table S1**). The identification of further three patients was facilitated by GeneMatcher (Sobreira *et al*., 2015). Genetic screening revealed homozygous missense variant in a second family (Family B): c.827C>A; p.Ala276Glu, while probands from Family C and D, from the same geographical area, had an in-frame deletion of 9 amino acids: c.346_372del; p.Glu116_Val124del (**Figure 1C**). These variants have not been previously described in patients; population frequencies are described in **Table S2**.

**Figure 1.**
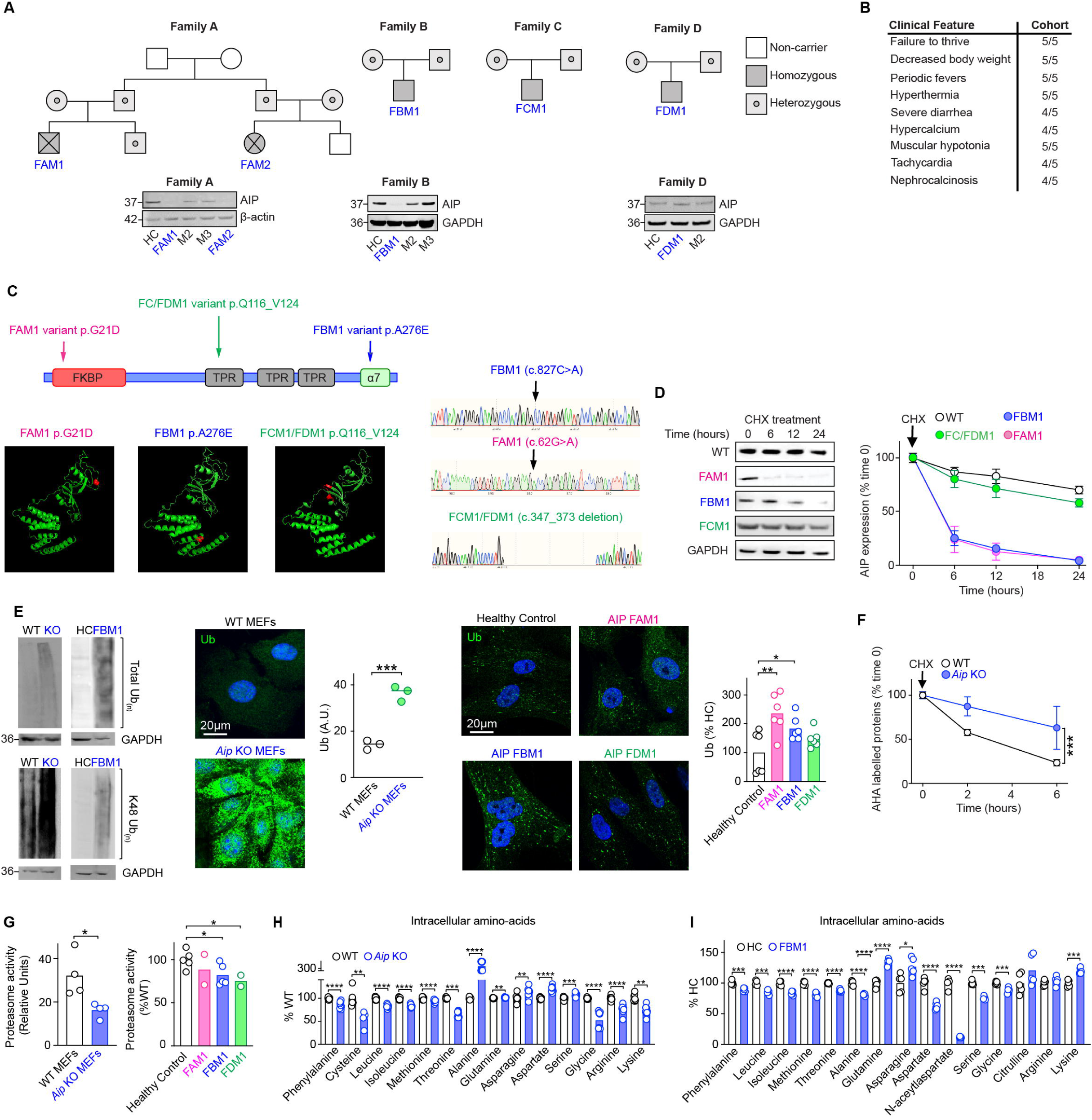
Family pedigrees of the four families of the children identified to have homozygous AIP variants and western blot analysis of AIP expression from patient (AIPd-PDFs) and healthy control (HC) derived dermal fibroblasts (**A**). Salient clinical features found in children born with AIP deficiency (**B**). Location of the variants in AIP (**C**). Family A and Family B AIP variants have a significantly reduced half-life as determined by cycloheximide (CHX) treatment of HEK cells transfected with plasmids containing *AIP* variants and analyzed by western blotting. Expression of AIP was normalized to a loading control (GAPDH) and to expression at time 0 Graphs show the mean ± SEM from at least two independent experiments (**D**). Expression of total ubiquitin and K48 ubiquitin from WT and *Aip* KO MEFs and from HC and AIPd-PDFs examined by western blotting and by immunofluorescence, arbitrary units (A.U) (**E**). WT and *Aip* KO MEFs were cultured with the methionine analogue L-Azidohomoalanine (AHA), treated with CHX and the relative amount of AHA determined by confocal microscopy (**F**). Proteasome activity was measured in WT and *Aip* KO MEFs and HC and AIPd-PDSFs (**G**). Levels of total amino acids determined by mass-spec analysis from WT and *Aip* KO MEFs (**H**) and from HC and PDF (**I**). Students un-paired *t*-test and 2-way ANOVA were used for analysis.

Analysis of AIP protein turnover analysis revealed that two of the three variants (p.Gly21Asp and p.Ala276Glu) resulted in proteins with significantly reduced half-life compared to wild type (WT) AIP (>24 hours), demonstrating that in these patients AIP was rapidly degraded as it is non-functional. The in-frame deletion variant showed a normal half-life (**Figure 1D**).

### AIP supports proteasome function

Loss of AIP in *Aip* KO MEFs and AIPd-PDSFs resulted in an increase in the total amount of ubiquitylated proteins (**Figure 1E**). We examined the rate of protein degradation incubating cells with a labelled methionine analogue, L-azidohomoalanine (AHA) that revealed that *Aip* KO MEFs had a reduction in the rate of protein degradation compared to WT MEFs (**Figure 1F**), leading us to examine the protein recycling mechanisms in cells lacking AIP. *Aip* KO MEFs and AIPd-PDFs displayed significantly less proteasome activity compared to WT cells (**Figure 1G**). *Aip* KO MEFs and AIPd-PDFs had significantly lower amounts of many amino acids compared to WT MEFs (**Figure 1H-I**). Autophagy can support the degradation of proteins when proteasome function is compromised (Kirkin *et al*., 2009; Suraweera *et al*., 2012; Korolchuk, Menzies and Rubinsztein, 2010), we therefore investigated if autophagy was impaired in *Aip* KO MEFs.

### AIP supports the initiation of autophagy

*Aip* KO MEFs failed to increase LC3 II expression upon starvation (with Earle’s Balances Salt Solution (EBSS)) or rapamycin treatment (**Figure 2A-C**). Confocal image analysis showed more ubiquitin–LC3 co-localization in *Aip* KO MEFs compared to WT MEFs, indicating that the ubiquitylated proteins that accumulated, were not being degraded by autophagy (**Figure S2A**). The accumulation of autophagosomes could not be induced by treating the MEFs with bafilomycin A1 (inhibits autophagosome-lysosome fusion), indicating that the defect in autophagy was due to the initiation of starvation-induced autophagy (**Figure 2B-C**). *Aip* KO MEFs had increased loss of viability when cultured in EBSS, EBSS plus bafilomycin and complete media plus bafilomycin compared to WT MEFs (**Figure S2B**), indicating that *Aip* KO MEFs were more sensitive to amino acid deprivation and dependent upon autophagy even under nutrient-rich conditions. To confirm that loss of AIP was responsible for the phenotype we observed, we lentivirus transduced human AIP in *Aip* KO MEFs to create a “rescue” cell line. *Aip* KO rescue MEFs had reduced expression of ubiquitin and Wipi2 and could induce LC3 (**Figure S2C-D**).

**Figure 2.**
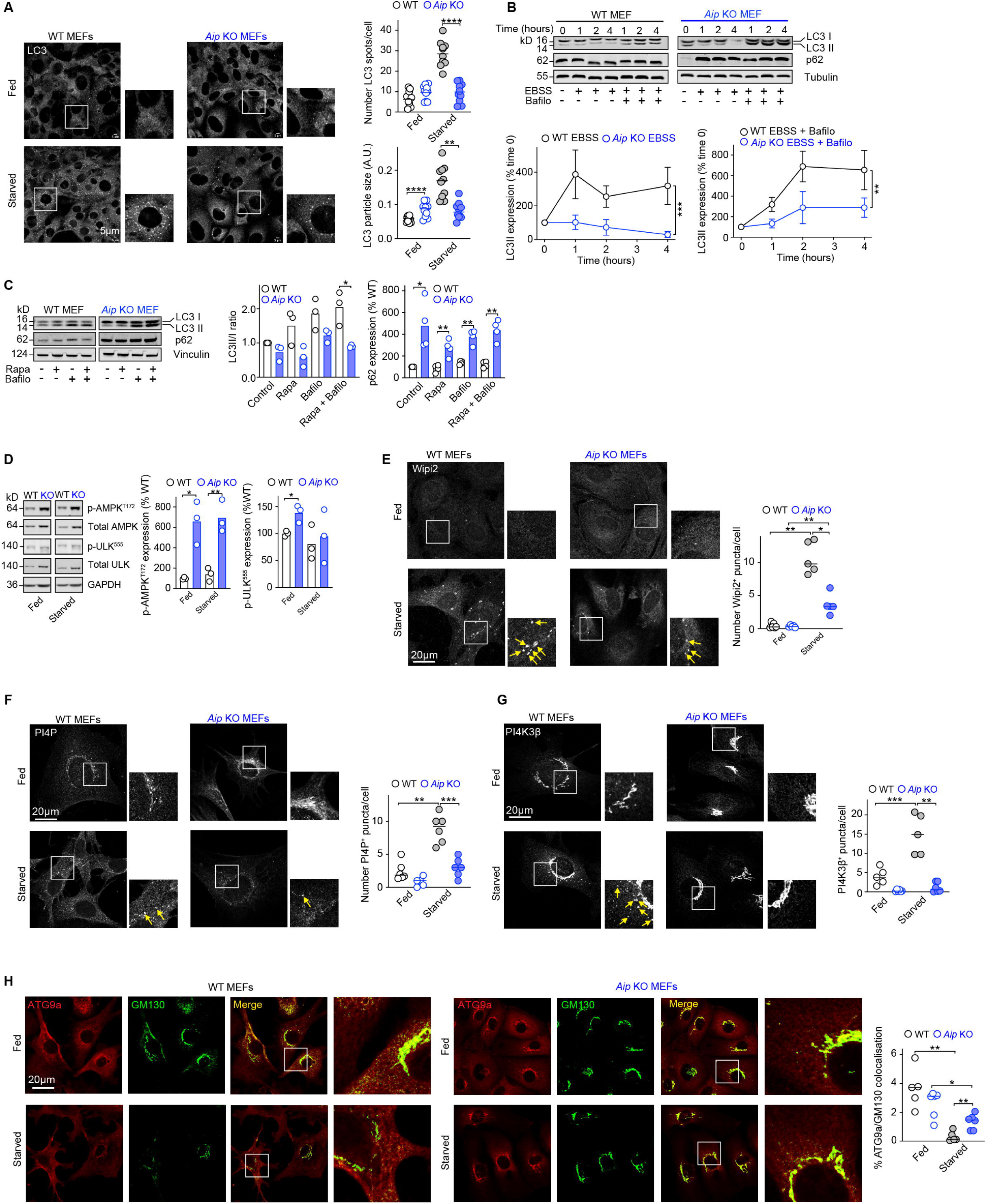
WT and *Aip* KO MEFs under fed and starved conditions (2 hours in Earle’s Balanced Salt Solution (EBSS)) and stained for LC3. The size and number of LC3^+^ puncta were determined (**A**). WT and *Aip* KO MEFs were cultured in EBSS (2 hours) in the presence or absence of bafilomycin A1 (Bafilo) (100nM) for 0, 2 and 4 hours and the expression of LC3I/II and autophagic flux determined (**B**). WT and *Aip* KO MEFs were treated with rapamycin (Rapa) (100nM) and Bafilo (100nM) for 2 hours and the expression of LC3I/II and p62 determined (**C**). Expression of p-AMPK, p-ULK^555^ in WT and *Aip* KO MEFs as determined by Western blotting (**D**). Wipi2^+^ puncta in WT and *Aip* KO MEFs under fed and starved (2 hours EBSS) conditions (**E**). Expression of PI4P^+^ (**F**), PI4K3β^+^ puncta (**G**), ATG9a/GM130 co-localization (**H**) in WT and *Aip* KO MEFs under fed and starved (2 hours EBSS) conditions. Graphs show the mean ± SEM from at least two independent experiments. Students un-paired and paired *t*-test and 2-way ANOVA used for analysis.

Nutrient deprivation activates AMPK, which in turn phosphorylates ULK1 to initiate autophagy (Gonzalez *et al*., 2020). *Aip* KO MEFs displayed signs of starvation under fed conditions; AMPK and the downstream autophagy activator p-ULK^S555^ were both activated under starved conditions, indicating that there was no defect in the upstream signaling events that activate autophagy in KO cells (**Figure 2D**). Phosphatidylinositol-3-phosphate (PI3P) formation on autophagy initiation sites (omegasomes) is required for the subsequent recruitment of ATG proteins (Dooley *et al*., 2014). PI3P was present in both *Aip* KO MEFs under fed and starved conditions in *Aip* KO MEFs, although less than WT MEFs under fed conditions, PI3P was still relatively abundant indicating that PI3P was unlikely to be the limiting factor in the initiation of autophagy in *Aip* KO MEFs (**Figure S2E**). Wipi2 is recruited to PI3P on omegasomes (Dooley *et al*., 2014); *Aip* KO MEFs had less Wipi2^+^ puncta/cell compared to WT MEFs under starved conditions, indicating a defect in the initiation of autophagy in *Aip* KO MEFs (**Figure 2E**).

Phosphatidylinositol-4-phosphate (PI4P) is produced at autophagosome initiation sites by the PI4K3β and PI4K2α enzymes recruited by ATG9a^+^ vesicles (Judith *et al*., 2019). This process precedes local PI3P production and has been shown to contribute to autophagosome recycling (McGrath *et al*., 2021; Judith *et al*., 2019). Under amino acid starvation, *Aip* KO MEFs showed a reduced number of PI4P^+^ puncta compared to WT MEFs (**Figure 2F**) and PI4P^+^ Wipi2^+^ puncta in response to starvation (**Figure S2F**), indicating that the altered generation and distribution of PI4P upon starvation might explain why *Aip* KO MEFs failed to fully initiate autophagy. Under fed conditions, PI4K3β had a peri-nuclear distribution, but upon starvation, distinct puncta were found throughout the cytoplasm in WT MEFs. In *Aip* KO MEFs, PI4K3β was located adjacent to the nucleus under fed and starved conditions in contrast to WT MEFs (**Figure 2G**), suggesting that there was a defect in the trafficking of PI4K3β to autophagy initiation sites (**Figure S2G**). ATG9a^+^ vesicles traffic from the Golgi to autophagy initiation sites (Judith *et al*., 2019). In *Aip* KO MEFs, more ATG9a was retained in the Golgi compared to WT MEFs (**Figure 2H**).

The transcription factors TFEB and TFE3 are crucial for the transcription of genes involved in autophagy migrating to the nucleus in response to starvation (Settembre *et al*., 2011). Under fed conditions, we observed increased expression of TFEB and nuclear TFE3 protein expression in *Aip* KO MEFs (**Figure S3A-B**). Moreover, key autophagy biogenesis genes were significantly up-regulated in *Aip* KO MEFs (**Figure S3C-E**), including *Atg9b*, *Ctsl1* and *Tfeb* under fed conditions (**Figure S3D-E**). In contrast, upon starvation, the expression of these genes did not increase in *Aip* KO MEFs with the exception of *Mcoln1* (**Figure S3D**). AIPd-PDFs had a similar higher baseline autophagy gene expression and inability to induce autophagy following starvation and similarly had many autophagy/lysosome genes expressed under fed conditions that failed to compensate for the defect in autophagy (**Figure S4A-D**).

### AIP supports lysosome function

Electron microscopic analysis revealed more autophagosomes and a decreased number of lysosomes in *Aip* KO MEFs and AIPd-PDF (**Figure 3A** and **Figure S4E**). A striking feature of electron microscopy analysis was an increase in the number and size of endosomes in *Aip* KO MEFs compared to WT MEFs (**Figure 3A**).

**Figure 3.**
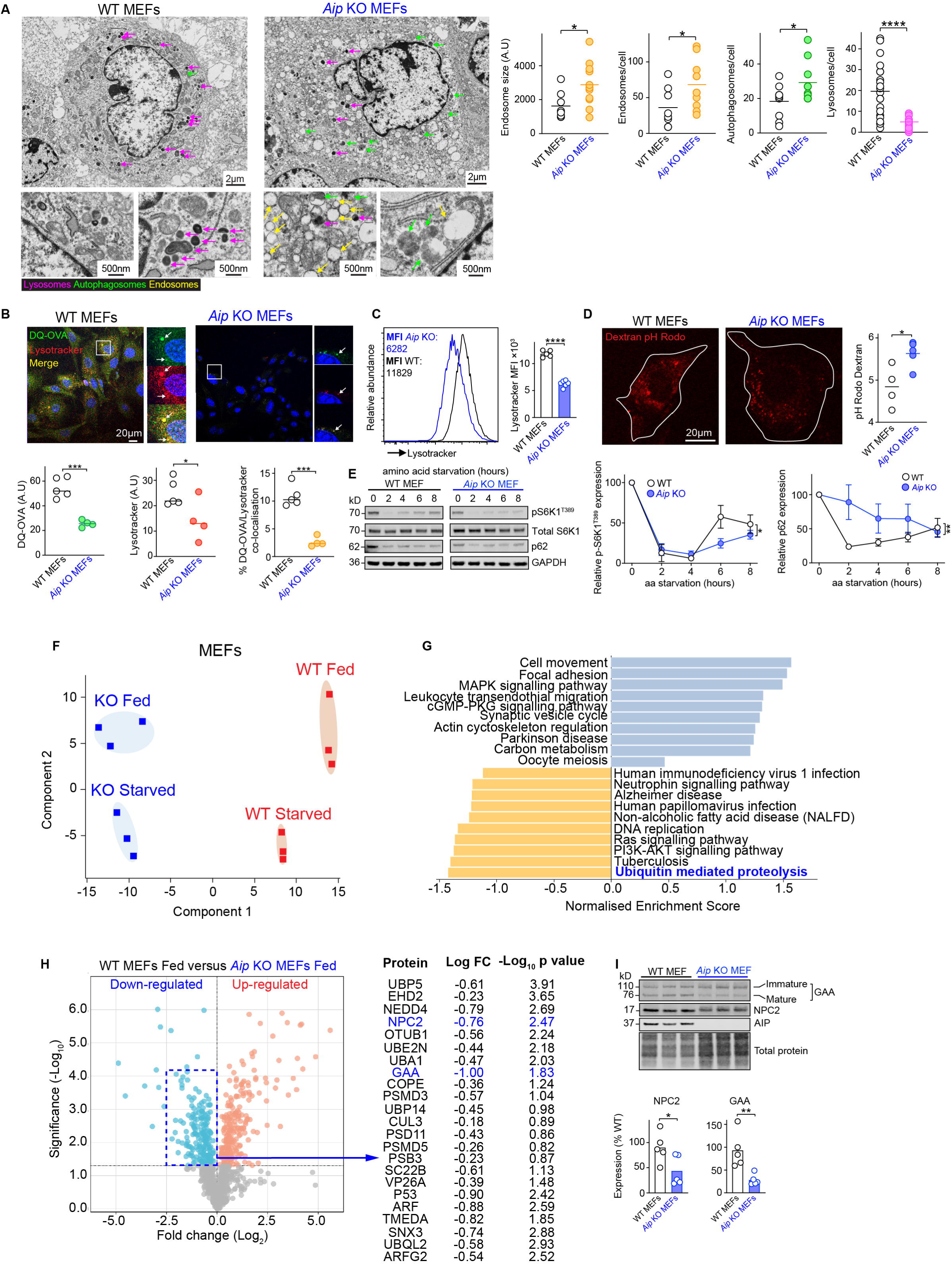
Electron microscopy of WT and *Aip* KO MEFs, highlighting the autophagosomes (green arrows), endosomes (yellow) and lysosomes (purple) (**A**). WT and *Aip* KO MEFs were incubated with DQ-OVA and stained with lysotracker (**B**). Expression of lysotracker in WT and *Aip* KO MEFs determined by flow cytometry (**C**). pH of endosomes determined by dextran-pH Rodo in WT and *Aip* KO MEFs (**D**). WT and *Aip* KO MEFs were starved of amino acids (EBSS media supplemented with 1% dialyzed FBS) and mTORC1 reactivation over time (pS6K1 expression) and p62 expression determined by western blotting (**E**). Principle component analysis of WT and *Aip* KO MEFs under fed and starved (2 hours EBSS) treatment (**F**). KEGG pathways analysis of TMT-mass spec analysis of WT versus *Aip* KO MEFs (**G**). Volcano plot showing significantly expressed protein between WT and *Aip* KO MEFs under fed conditions (**H**). Expression of NPC2 and GAA in WT and *Aip* KO MEFs normalized to total protein (**I**). Graphs show the mean ± SEM from at least two independent experiments. Students un-paired, paired *t*-test and 2-way ANOVA were used for analysis.

AIP-deficient cells had an inability to digest DQ-ovalbumin (OVA) and showed decreased lysotracker expression, suggesting reduced lysosome function (**Figure 3B-C**). AIPd-PDSFs showed a similar defect (**Figure S4F**). RNA-sequencing analysis revealed that *Aip* KO MEFs had reduced expression of many lysosome genes (**Figure S5A**) and reduced expressed of many genes associated with lysosomal storage diseases (LSDs) (**Figure S5B**). The pH of the lysosomes was higher in *Aip* KO MEFs (∼pH 5.6) than WT MEFs (∼pH 4.4) demonstrating reduced lysosome acidification (**Figure 3D**).

### AIP is required for autophagic lysosome reformation

The inability to continue autophagy during amino acid starvation indicated that there might be a defect in autophagosome recycling. Upon the induction of autophagy, mTORC1 is reactivated following the liberation of amino acids from lysosomes. *Aip* KO MEFs had a delay in the reactivation of mTORC1 and a delay in p62 degradation indicating a defect in autophagic lysosome reformation (ALR) (Yu *et al*., 2010) (**Figure 3E**). Confocal analysis of lysosomes in *Aip* KO MEFs showed enlarged lysosomes at 4 hours starvation before becoming smaller, in accordance with a defect in ALR (**Figure S5C**). Feeding *Aip* KO MEFs with albumin following starvation also resulted in a delay of mTORC1 activation (**Figure S5D**). mTORC1 activation in response to leucine has been shown to be dependent upon functional lysosomes (Wyant *et al*., 2017). *Aip* KO MEFs had a muted response to leucine in contrast to WT MEFs, but had a robust response to glutamine, which is not lysosome dependent (Jewell *et al*., 2015), further supporting a role for impaired lysosome function in *Aip* KO MEFs (**Figure S5E**).

As AIP is a promiscuous co-chaperone molecule and based upon the observation that loss of AIP results in an increase in ubiquitylated proteins, we sought to gain insight into how AIP contributed to the loss of proteostasis. We performed unbiased Tandem Mass Tagging (TMT) analysis of WT and *Aip* KO MEFs. Principle component analysis of the proteomic data revealed clear differences between WT and *Aip* KO MEFs under fed and starved conditions (**Figure 3F**) and that the most down-regulated process under fed conditions was ubiquitin mediated proteolysis supporting our original observation (**Figure 3G**). Proteins with reduced expression in *Aip* KO MEFs included those involved in ubiquitin-mediated proteolysis, proteasome subunits, vesicle trafficking and lysosomal function, such as alpha glucosidase (GAA) and Niemann-Pick disease type C Protein 2 (NPC2) (**Figure 3H-I**).

### Loss of AIP results in metabolic reprogramming

Despite the reduced protein recycling in cells lacking AIP, cells grew well *in vitro* and were outwardly normal, indicating that AIP deficient cells were adapting to the shortage of amino acids. *Aip* KO MEFs underwent increased glycolysis, importing more glucose, producing less ATP and more reduced NADH than WT MEFs (**Figure 4A-E**). Metabolomic analysis of *Aip* KO MEFs revealed that they had increased total amounts of glycolysis products (pyruvate, lactate, alanine) and increased proportion of glucose isotopologues derived from uniformly labelled glucose (U-^13^C_6_) (m+3) compared to WT MEFs (**Figure 4F**). Similar results were obtained from the analysis of supernatant of cultured *Aip* KO MEFs (**Figure S6A-B**). *Aip* KO MEFs and AIPd-PDSFs were more sensitive to glucose deprivation (**Figure S6D-E**) and glycolysis inhibition by oligomycin and 2-deoxy-d-glucose (2-DG) (**Figure S6F**). *Aip* KO MEFs had a greater proportion of TCA cycle intermediates including malate, α-ketoglutarate (α-KG), aspartate and aconitate derived from labelled glucose (m+3) (**Figure S7A-C**). We found that *Aip* KO MEFs incorporated more labelled glucose (m+3) into glucogenic amino acids including alanine, serine, aspartate, asparagine, glutamate and aspartate compared to WT MEFs (**Figure S7D**) and had increased expression of many genes and proteins involved in oxidative phosphorylation (**Figure S7E-F**). *Aip* KO MEFs had increased mitochondrial membrane potential, mitochondrial volume and ROS production (**Figure 4G-J**). Interestingly, we observed that the increased glycolysis in *Aip* KO MEFs was being utilized for serine-1 metabolism (Yang and Vousden, 2016) (**Figure S7G-I**). Together, these results provided evidence of increased aerobic glycolysis and TCA cycle activity in cells lacking AIP.

**Figure 4.**
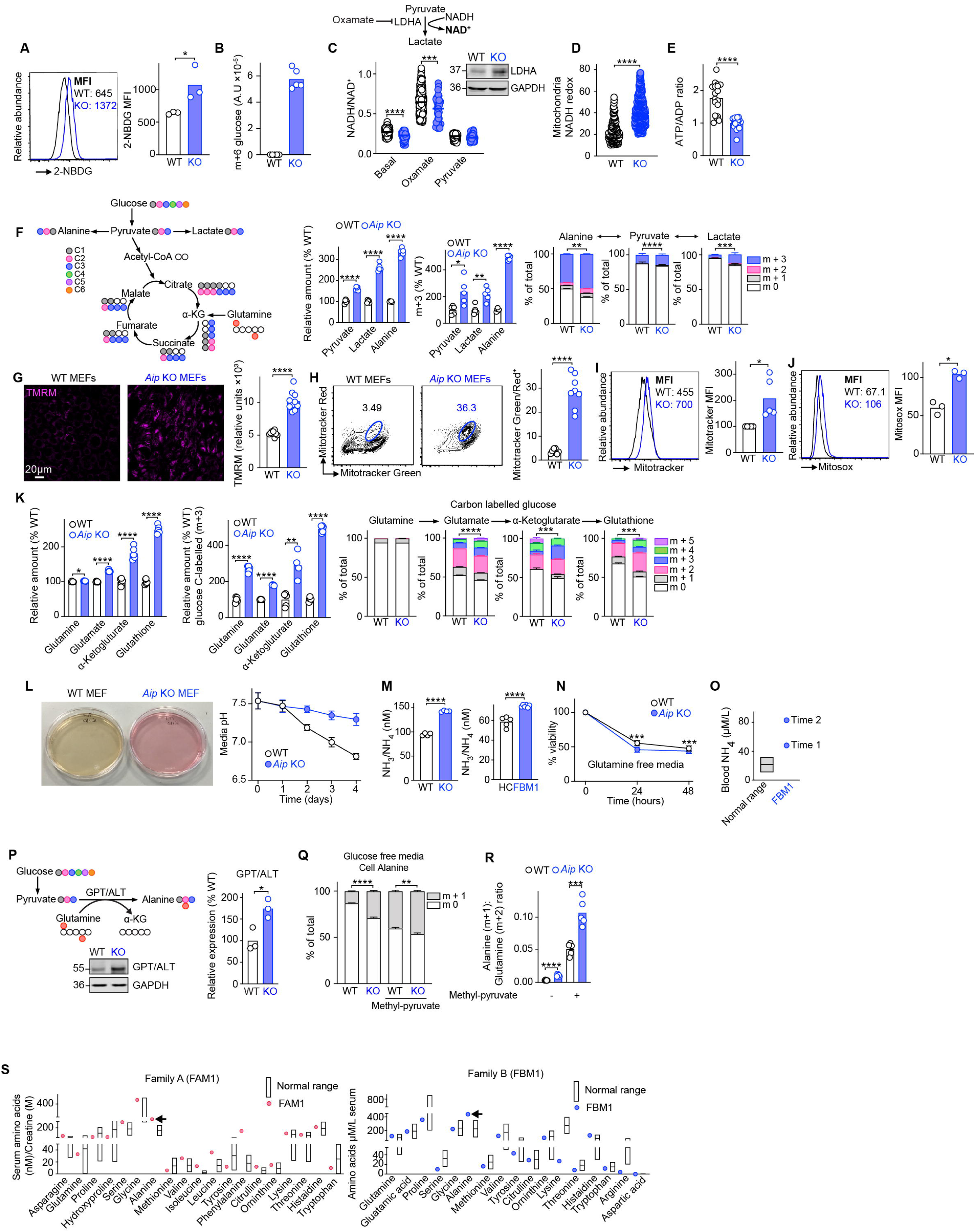
Glucose import into WT and *Aip* KO MEFs was determined using the fluorescent glucose analogue 2-NBDG (**A**) and the relative amount of uniformly labelled glucose (U-^13^C_6_ [m+6]) in WT and *Aip* KO MEFs (**B**). The rate of glycolysis was measured by treating the cells with sodium oxamate and measuring the ratio of NAD/NADH in WT and *Aip* KO MEFs (**C**) Mitochondrial NAD/NADH was measured in WT and *Aip* KO MEFs (**D**). The ATP/ADP ratio was determined in WT and *Aip* KO MEFs (**E**). Metabolomic analysis using uniformly labelled glucose (U-^13^C_6_) in WT and *Aip* KO MEFs measuring total glycolysis products and flux (m+3) from labelled glucose (**F**). Mitochondrial membrane potential was determined in WT and *Aip* KO MEFs using Tetramethylrhodamine methyl ester (TMRM) (**G**). Mitochondria volume and membrane potential was determined in WT and *Aip* KO MEFs using Mitotracker red/Mitotracker green (**H**), mitotracker (**I**) and mitosox (**J**) dyes by flow cytometry and median fluorescent intensity (MFI) determined. Relative levels of glutamine metabolites in WT and *Aip* KO MEFs and relative levels of glutamine metabolites and flux derived from uniformly labelled (U-^13^C_6_) glucose in WT and *Aip* KO MEFs (**K**). Media pH over time in WT and *Aip* KO MEFs (**L**). Levels of NH_3_/NH_4_ in WT and *Aip* KO MEFs and HC and PDSFs (**M**). Viability of WT and *Aip* KO MEFs cultured in glutamine free media (**N**). Elevated blood NH_4_ levels determined from patient F538M1 compared to the normal range (**O**). Transamination of glutamine using pyruvate as an amino acceptor by the enzyme GPT/ALT expressed in WT and *Aip* KO MEFs (**P**). WT and *Aip* KO MEFs were cultured in glucose free media with U-^15^N_2_ glutamine in the presence or absence of 10mM methyl-pyruvate and the relative proportion of U-^15^N_2_ glutamine isotopologues determined (**Q**). The ratio of alanine (m+1): glutamine (m+2) (**R**). Serum amino acids from F487M1 and F538M1 were analyzed and compared against the normal range for these amino acids. Arrows highlight alanine (**S**). Graphs show the mean ± SEM from at least two independent experiments. Students un-paired, paired *t*-test and 1-way and 2-way ANOVA were used for analysis.

*Aip* KO MEFs had increased amounts of glutamine metabolites (**Figure 4K**) within cells and in the culture supernatant (**Figure S6E**). Culturing *Aip* KO MEFs, we noticed that the media pH remained high compared to WT MEFs (**Figure 4L**) and that *Aip* KO MEFs and AIPd-PDFs had increased levels of ammonia (NH_3_/NH_4_) compared to WT MEFs (**Figure 4M**). We hypothesized that ammonia released as a result of increased transamination contributed to maintaining the pH (**Figure S8A**). *Aip* KO MEFs had increased expression of solute channels that import glutamine into cells (*Slc1a5, Slc25a22, Slc38a1*) and altered expression of enzymes involved in glutaminolysis (**Figure S8B-C**). *Aip* KO MEFs more sensitive to glutaminolysis inhibitors than WT MEFs (**Figure S8D**). Under conditions of increased aerobic glycolysis, cells can maintain TCA cycle intermediates using anaplerosis, (Yang, Venneti and Nagrath, 2017; Zhang, Pavlova and Thompson, 2017). We hypothesized that the glutamine would be converted into α-ketoglutarate entering the TCA cycle, but examination of TCA cycle intermediates revealed that less intermediates were derived from U-^13^C_5_ glutamine in *Aip* KO MEFs and we observed neither an increase in oxidative or reductive glutamine metabolism in *Aip* KO MEFs (**Figure S8E**), but there were differences in the utilization of U-^13^C_5_ and U-^15^N_2_ glutamine in glutaminolysis metabolites and serine and alanine (**Figure S8F**). Interestingly, patient FBM1 had an increase amount of ammonia in blood with normal liver function compared to the normal range on two separate occasions, providing evidence that increased glutaminolysis was occurring (**Figure 4O**).

Transamination is an exchange of an amino group to another molecule and is important in the metabolism of amino acids. In starvation, amino acids are deaminated to provide TCA cycle substrates to reinforce energy generation and protect essential cellular functions (Torres *et al*., 2023). The increased glycolysis observed in *Aip* KO MEFs was being utilized for transamination as the resulting pyruvate would be used as an amino acceptor producing alanine. *Aip* KO MEFs had increased expression of the transaminase GPT/ALT (**Figure 4P**). Analysis of WT and *Aip* KO MEFs cultured in the absence of glucose supplemented with methyl-pyruvate demonstrated increased transamination of the amino group of U-^15^N_2_ glutamine onto pyruvate producing significantly more U-^15^N_1_ (m+1) labelled alanine with an increase in the ratio between (m+1) labelled alanine and (m+2) glutamine (**Figure 4Q-R**). Interestingly the serum from two patients showed increased levels of alanine (**Figure 4S**). Together, these results indicated that *Aip* KO MEFs had evidence of increased transamination, a mechanism by which AIP deficient cells use to relieve the toxicity of protein accumulation.

### AIP loss in zebrafish recapitulates patient and MEF phenotype

To gain a better understanding of the complex phenotype of the AIP deficient children, using CRISPR-Cas-9 we deleted *aip* in zebrafish (**Figure S9A-B**). Similar to the children born with homozygous loss of AIP, homozygous *aip* KO fish showed reduced growth rate and failure to thrive, being significantly smaller by day 6 post-fertilization (dpf) (**Figure 5A**). There was no difference in the survival between WT and *aip* KO fish between days 5-6 dpf. But the survival of *aip* KO fish markedly decreased afterwards, coinciding with depletion of the yolk (6 dpf) (**Figure 5B**, **Figure S9C**). In contrast to Aip KO MEFs and AIPd-PDSFs, *aip* KO fish had less ubiquitylated proteins, likely a result of reduced protein translation in response to chronic starvation (**Figure 5C**) and reduced level of some amino acids (**Figure 5D**), but still had reduced proteasome activity (**Figure 5E**) and increased ammonia production (**Figure 5F**). Autophagy has been shown to be essential for transition from the larval to juvenile stages of zebrafish development (Maweed Suzan Attia, 2021) and essential for neonatal survival in mice (Efeyan *et al*., 2013; Kuma *et al*., 2004; Komatsu *et al*., 2005). *aip* KO fish expressed less LC3 and failed to induce autophagy on dietary restriction (**Figure 5G**) and reduced lysosome function (less lysotracker expression) consistent with AIP deficient fibroblasts (**Figure 5H**).

**Figure 5.**
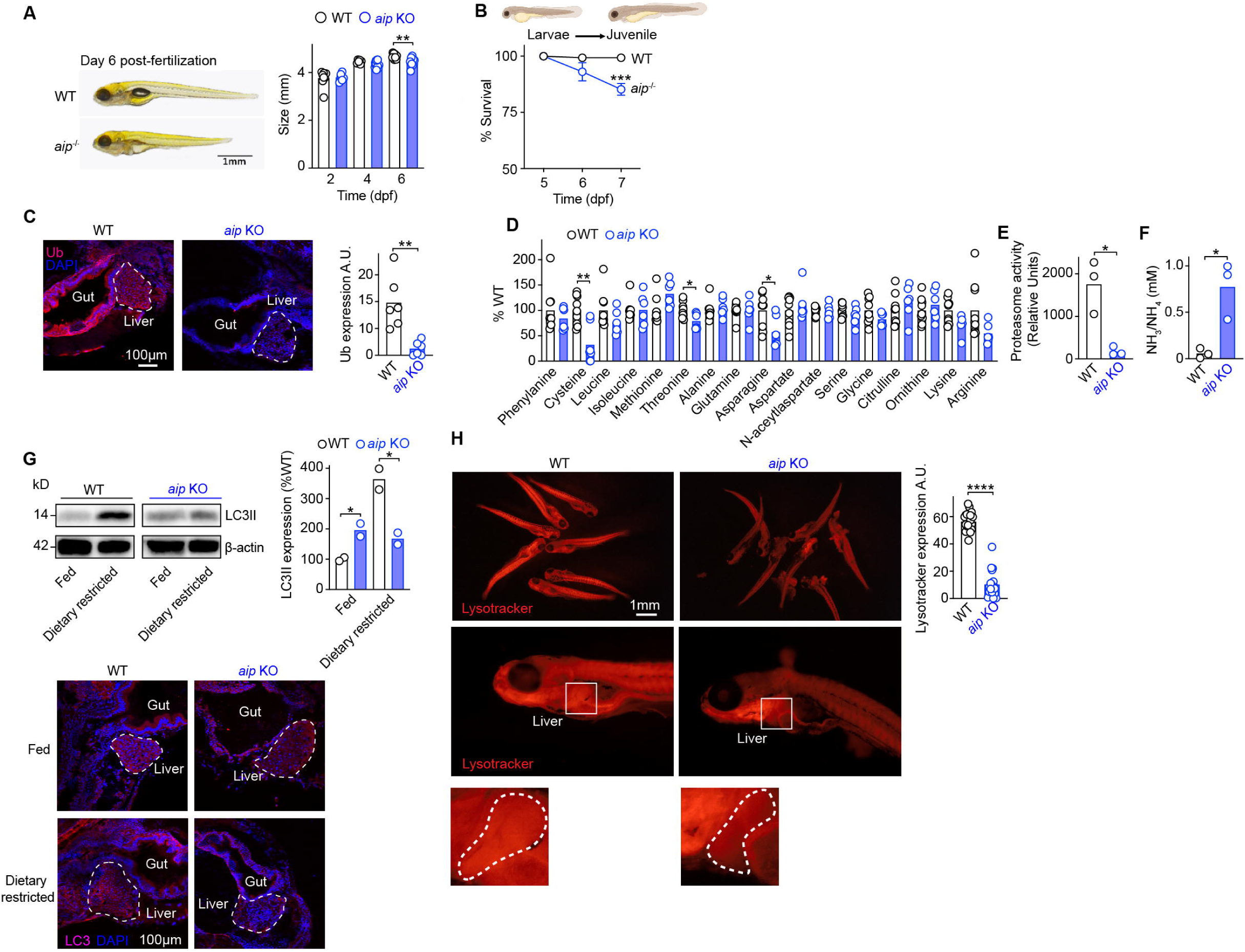
Size of WT and *aip* knock out zebrafish (KO) from day 2 dpf (**A**). Survival rate of WT and *aip* mutant zebrafish from day 5 post fertilization (dpf) under fed and dietary restricted conditions (**B**). Ubiquitin expression examined by immunofluorescence (**C**). Relative amino acids from WT and *aip* KO zebrafish (**D**). Relative proteasome activity in WT and *aip* KO zebrafish (**E**). Ammonia (NH_3_/NH_4_) concentration from WT and *aip* KO zebrafish (**F**). LC3 expression under fed and dietary restriction (**G**). Lysotracker expression in WT and *aip* KO zebrafish (**H**). Fish used for experiments were between 5-7 dpf. Graphs show the mean ± SEM from at least three independent experiments. Students un-paired and paired *t*-test used for analysis.

To determine if *aip* KO fish underwent metabolic reprogramming, we treated day 4 dpf fish with tracer metabolites (glucose U-^13^C_6_ and glutamine U-^13^C_5_). Similar to what we observed in *Aip* KO MEF, *aip* KO fish had evidence of increased glycolysis, TCA cycle activity and glutaminolysis (**Figure S10A-C**).

## Discussion

Children born with homozygous biallelic loss of function variants of AIP resulted in a severe metabolic disease which is refractory to current medical intervention. Understanding the molecular pathobiology of this novel disease is therefore imperative for the treatment of these children. We identified several potential mechanisms by which loss of AIP contributes to the disease pathology manifest in these children.

AIP deficient children were born with normal body size parameters but demonstrated growth retardation and an inability to gain weight despite being on total parenteral nutrition therapy indicating that defective nutrient absorption from the gut was not the major reason why they failed to gain weight. Using *aip* KO zebrafish, we observed a similar phenotype; viable until the yolk was depleted, subsequently dying following transition from the larvae to juvenile stage of development, when autophagy is essential (Maweed Suzan Attia, 2021). Patients deficient in ATG7, an essential autophagy protein have a milder phenotype than the AIP deficient patients (Collier *et al*., 2021). This is likely due to the impairment of several key UPS processes that AIP supports in addition to the initiation of autophagy.

In all three species in which AIP was deficient, metabolic reprogramming occurred. Excessive protein catabolism in *Aip* KO MEFs produces more ammonia, neutralizing the natural pH decline of the media due to the accumulation of acidic lactate. This scenario is supported by a reduction in both intra and extracellular amino acid levels in *Aip* KO MEFs (except those acting as amino acceptors) and enhanced transamination of glutamine to alanine. Increased circulating ammonia levels and a tendency for reduced blood amino acid levels have been confirmed in AIP deficient patients. The corresponding human, murine and zebrafish data suggest that the amino acid catabolism that exceeds the body’s capacity (via the urea cycle) to detoxify it. In parallel, with the major protein catabolic routes blocked and vulnerability to energy deprivation, AIP deficient cells may switch to an energy-sparing survival mode. The reduction of energy requirements may not be a workable option for cells with constitutively high energy demands that occurs during post-natal development.

The description of a novel metabolic disease caused by homozygous loss of the *AIP* gene provides important insights into the mechanisms that support proteostasis and metabolic processes. Chaperone molecules are some of the most abundant proteins within cells, but the precise role of many of these chaperone molecules is unknown (Saibil, 2013; Hartl, Bracher and Hayer-Hartl, 2011). The striking phenotype we observed in AIP deficient children, cells and zebrafish indicates that AIP supports proteostasis and key hubs of cellular metabolism. Understanding the mechanisms that support the initiation of autophagy, lysosome and the proteasome are of fundamental importance.

Lysosomal storage diseases (LSDs), are individually rare, but collectively relatively common group of diseases (Platt *et al*., 2018; Pechincha *et al*., 2022; Castellano *et al*., 2017; Parenti, Medina and Ballabio, 2021). This novel disease shares features of LSDs with evidence of reduced lysosome activity, impaired vesicle trafficking, reduced autophagic flux, an accumulation of autophagosomes, loss of calcium homeostasis, increased oxidative stress and reduced expression of many genes associated with LSDs, however the clinical and cellular phenotype observed makes it distinct from other LSDs.

In conclusion, we provide the first description of homozygous loss of AIP in humans that results in a severe multi-faceted metabolic disease revealing the many roles in which the chaperone molecule AIP supports metabolic processes within cells in particular supporting maintenance of proteostasis within cells (**Figure S11A-D**).

## Ethics

Parents signed written consent to take part in the study and the study was approved by the Ethic Committee (MREC 06/Q0104/133)

CB Home Office license number: PP9581087

## Competing interests

No competing interests.

## DATA and materials availability

MEFs and *aip* KO zebrafish used in this study are available upon request.

## Author contributions

MK obtained patient data and samples, conceived the studies along with OH, designed some experiments, wrote parts of the manuscript and obtained funding. SB, CCL, OS, NU, LV, performed experiments and helped analyze data throughout this study. SMH, VM, KB, MD, KS, CYC and GB helped design, perform and experiments for the metabolic analysis. CH helped analyze RNA-sequencing and proteomics data. LP and JPC helped with confocal microscopy and metabolic analysis. HTB, VÖE, KAR, HVE, VEK, AD, RZ and AB identified and cared for the patients, performed original genetic testing and provided clinical data. GA performed electron microscopy and helped analyze the images. SDT and ST performed experiments, analyzed data and provided intellectual support and reagents. EA provided intellectual support, helped design experiments and analyzed data. CB and XW generated *aip* mutant zebrafish and designed and performed zebrafish experiments. OH conceived the study, designed, performed and analyzed experiments and data, wrote the manuscript and obtained funding for this study.

## Acknowledgments

This study was supported by grants from the Rosetrees Trust [M789] and the Great Ormond Street Hospital Children’s Charity (registered charity no.1160024) [V4722]) to OH and MK. A. core confocal microscopy facility for the William Harvey Research Institute (MGU093) was awarded to J.P Chapple. CTL was supported by a Clinical Training Fellowship from The Medical College of St Bartholomew’s Trust, XW was supported by a PhD scholarship the China Scholarship Council. We are grateful for the patients and their families for providing samples and data for the study. We acknowledge the help of Dr. Steve Lynam and Dr. Xiaoping Yang from the Kings College London Proteomics facility. We are grateful to Dr. James Davison and Dr. Detlef Bockenhauer at Great Ormond Street Hospital in London, UK and to numerous colleagues who have provided reagents and helpful discussions regarding this work.

---

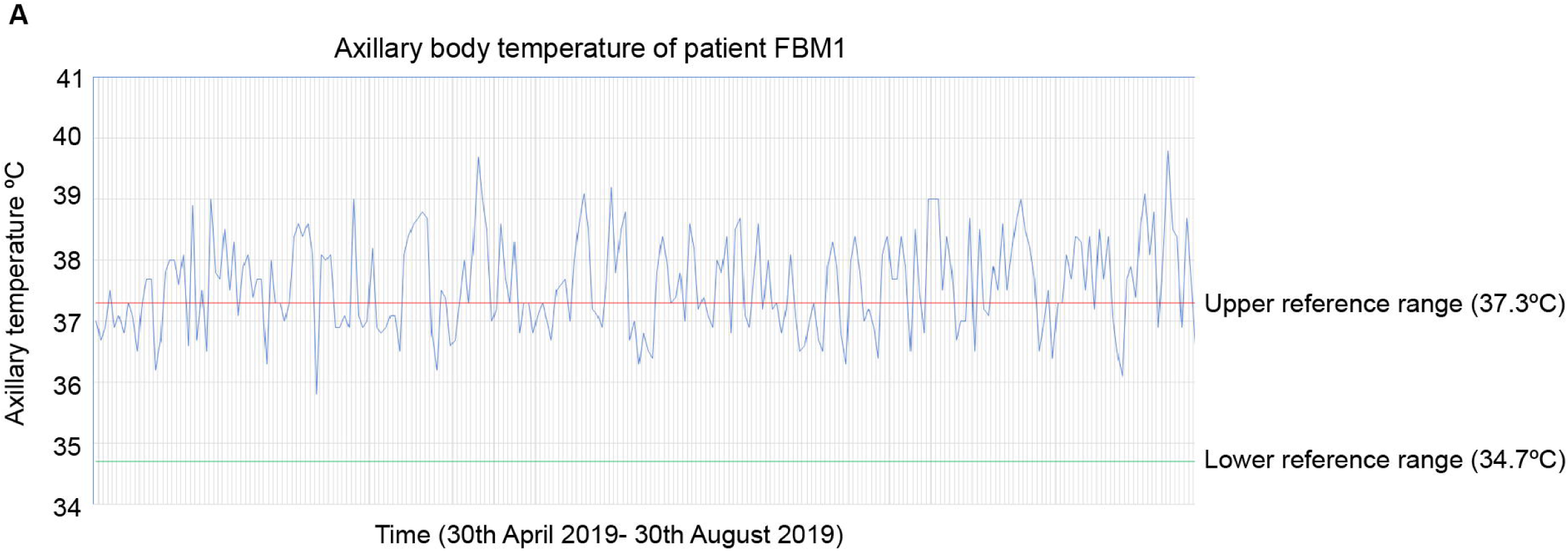
Figure S1

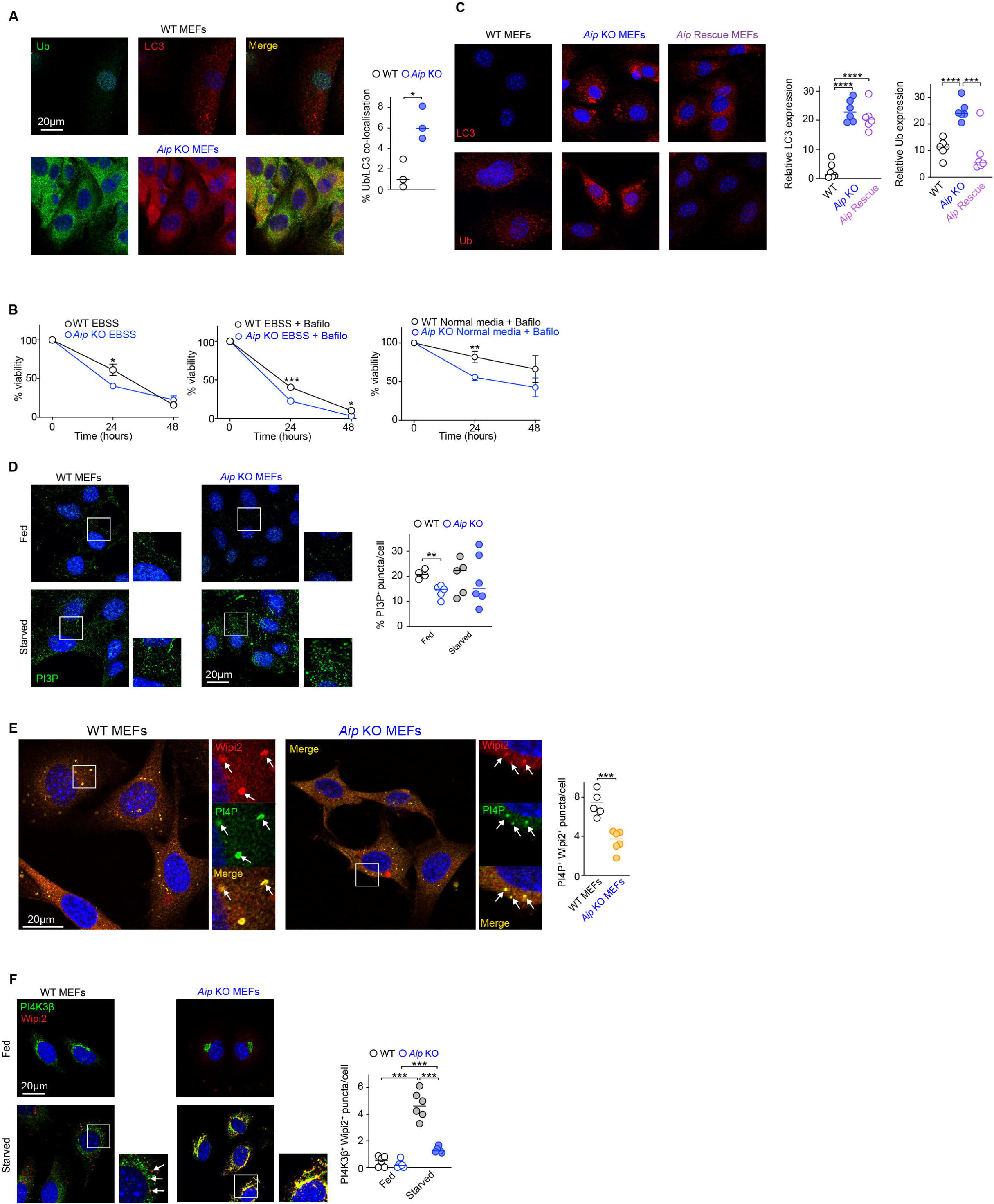
Figure S2

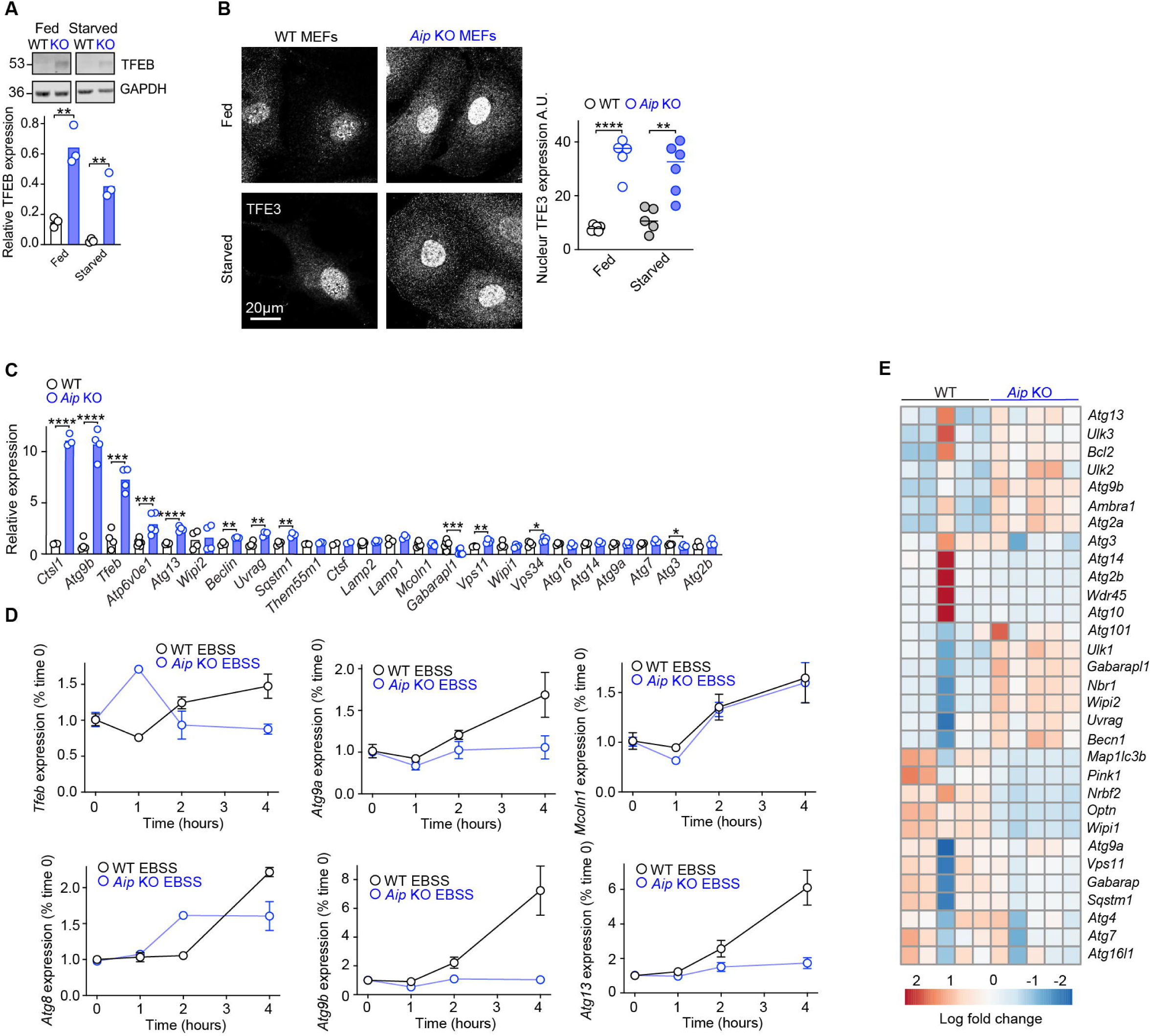
Figure S3

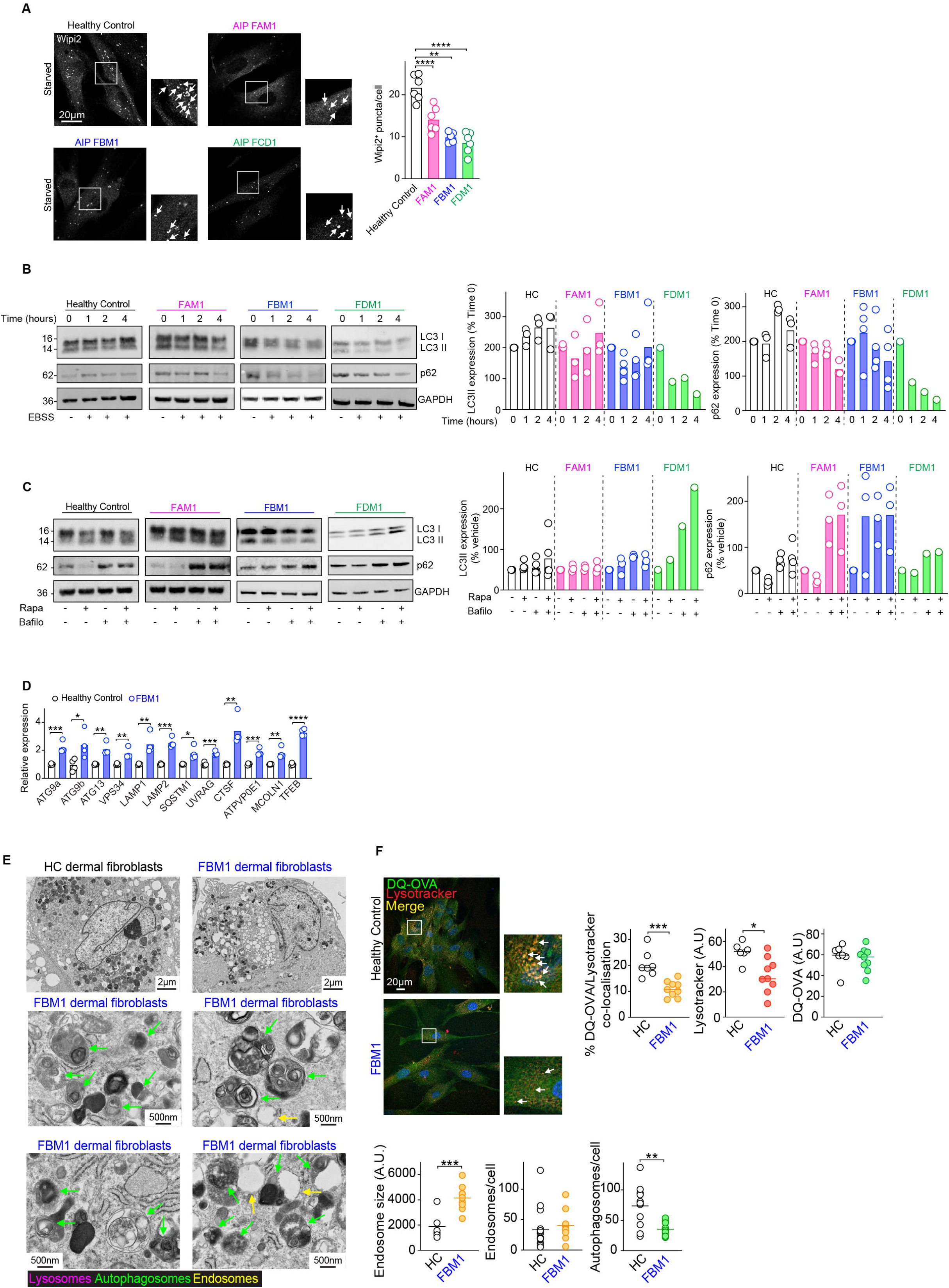
Figure S4

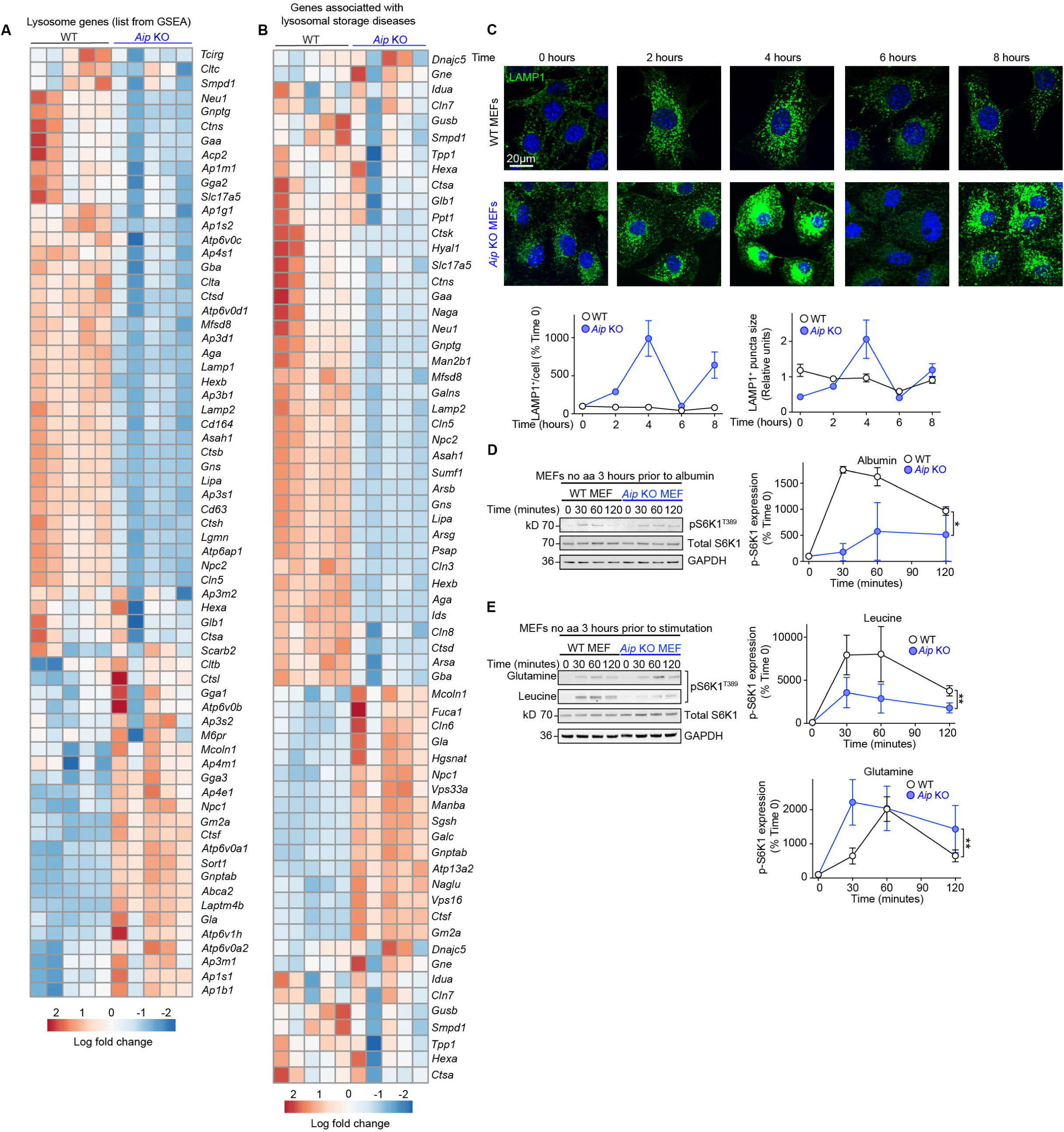
Figure S5

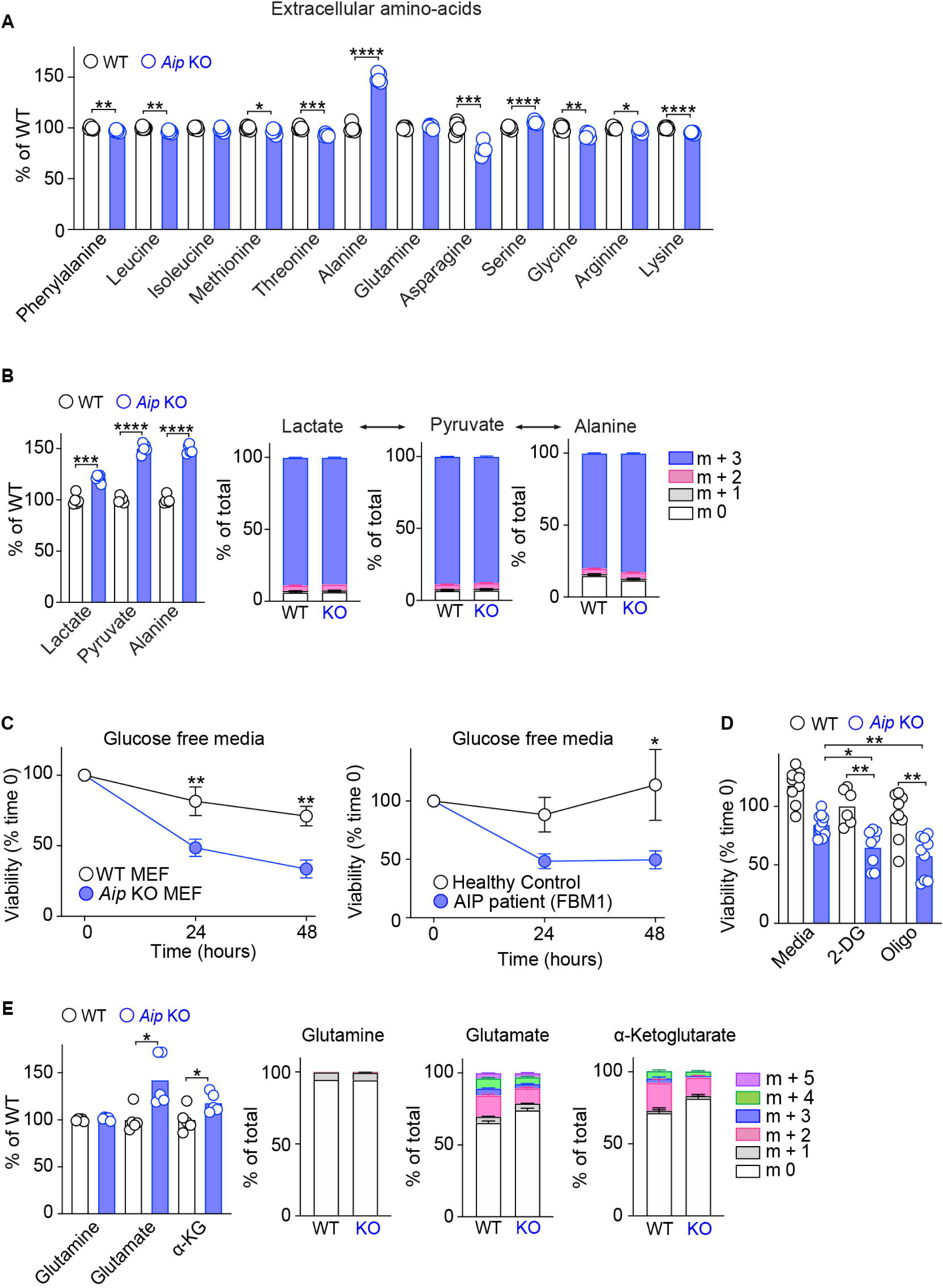
Figure S6

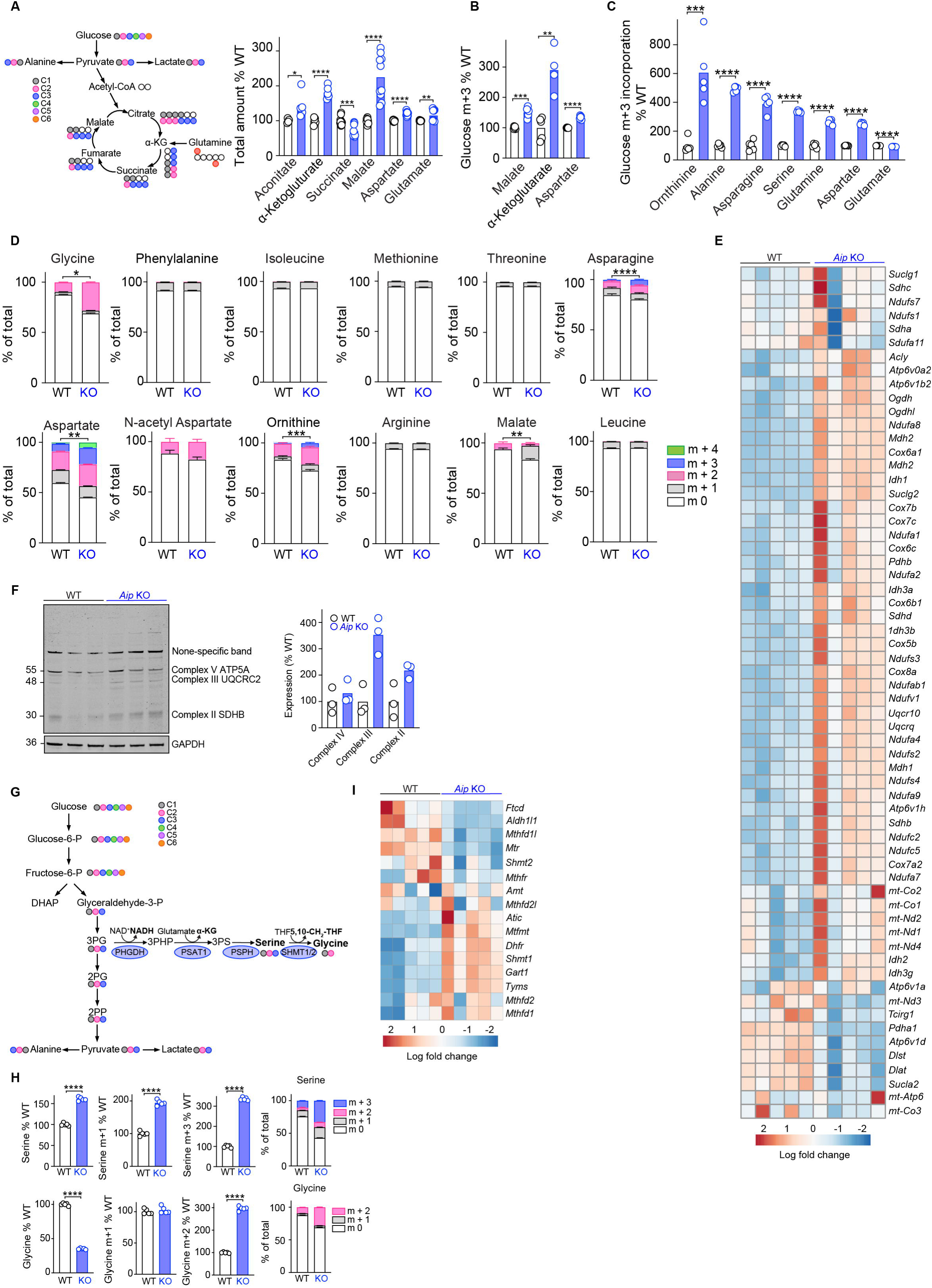
Figure S7

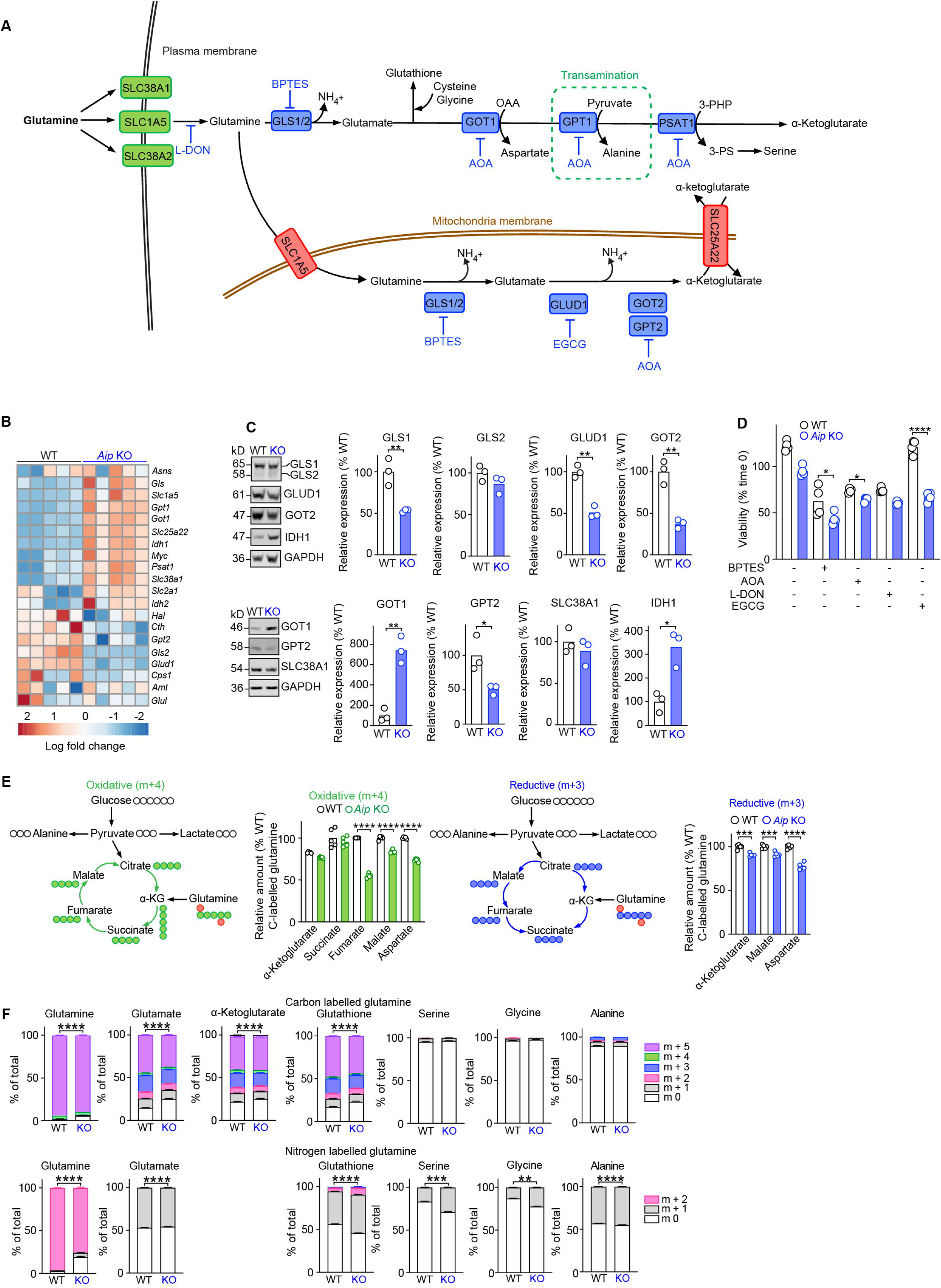
Figure S8

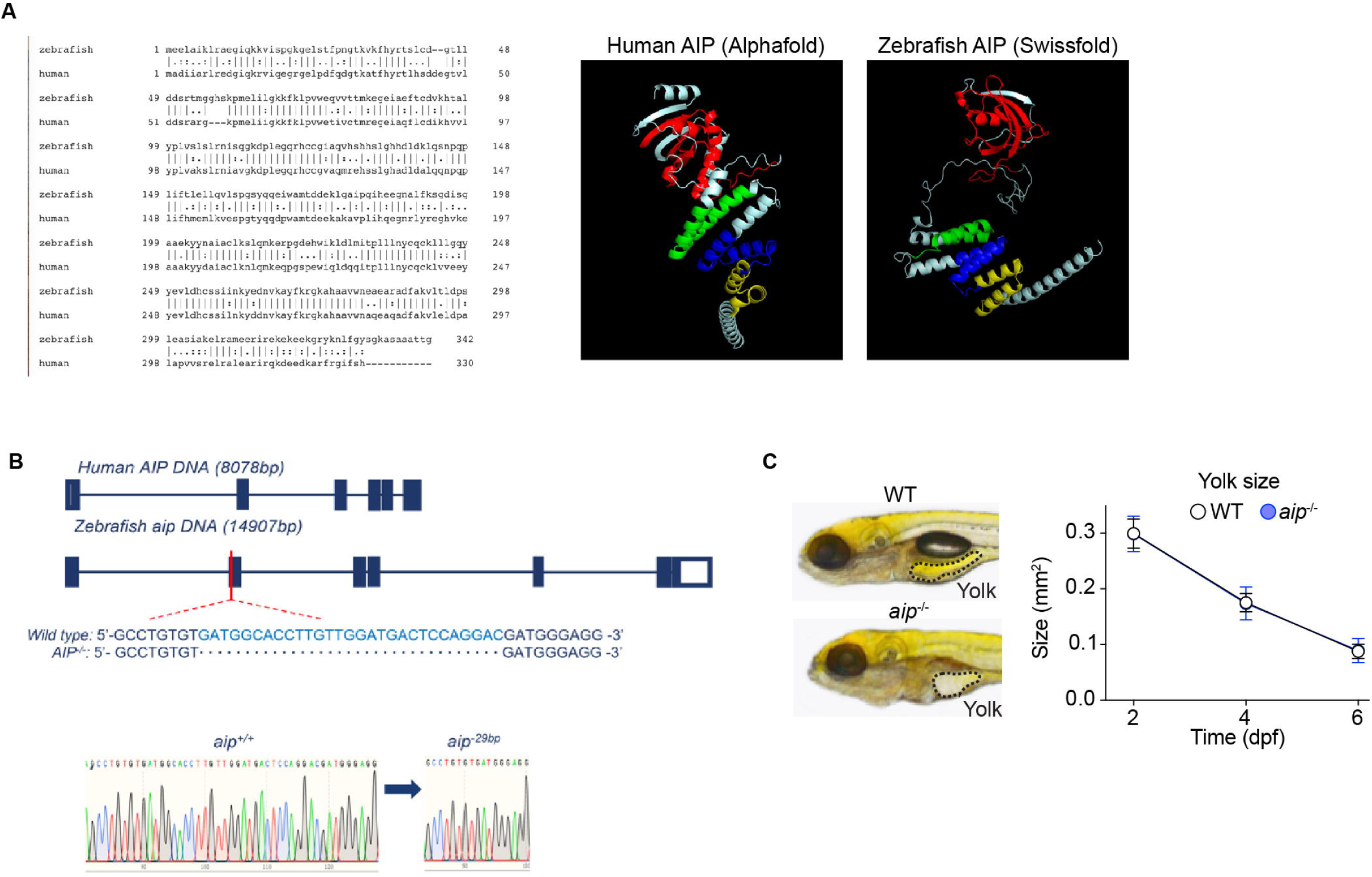
Figure S9

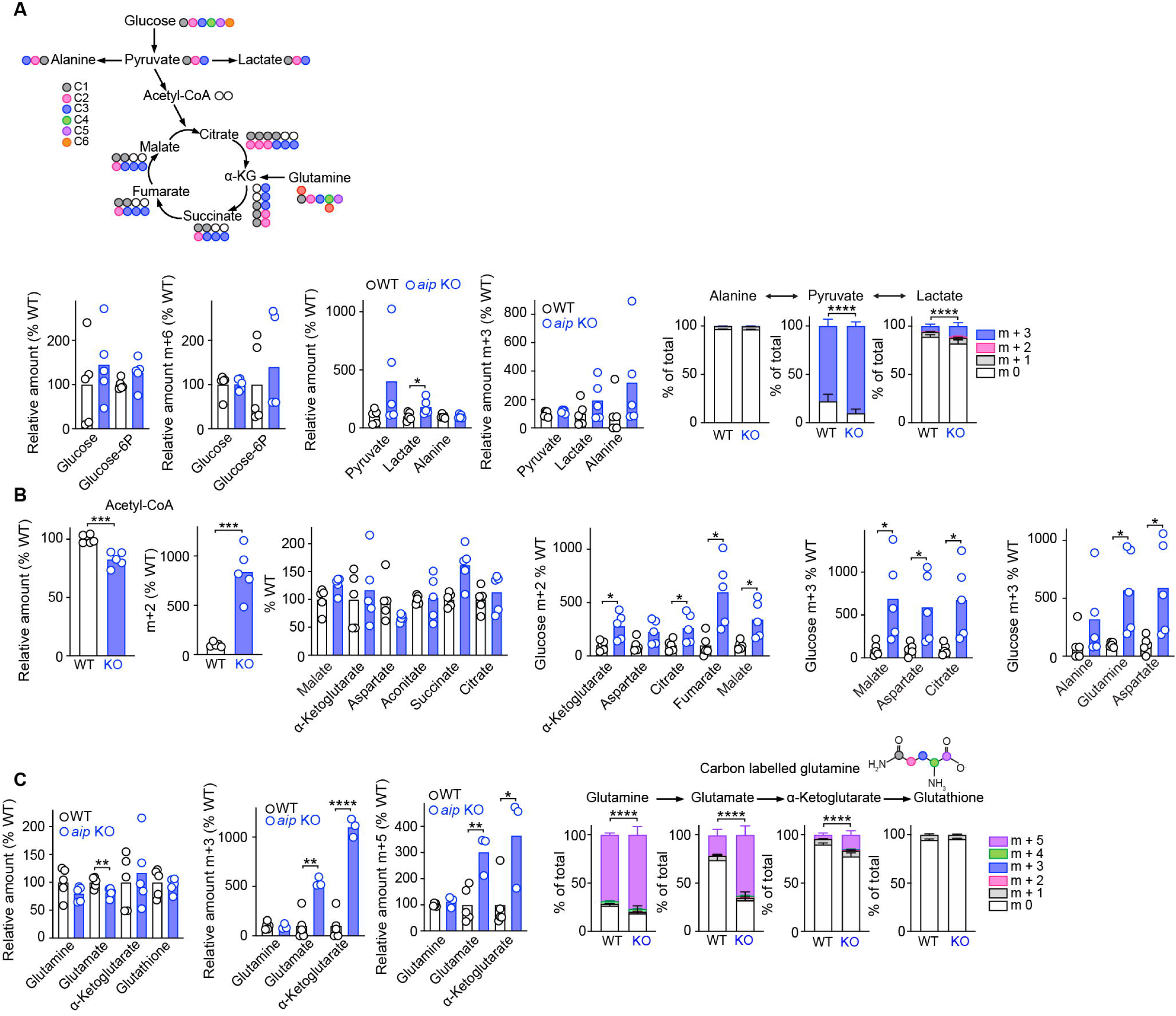
Figure S10

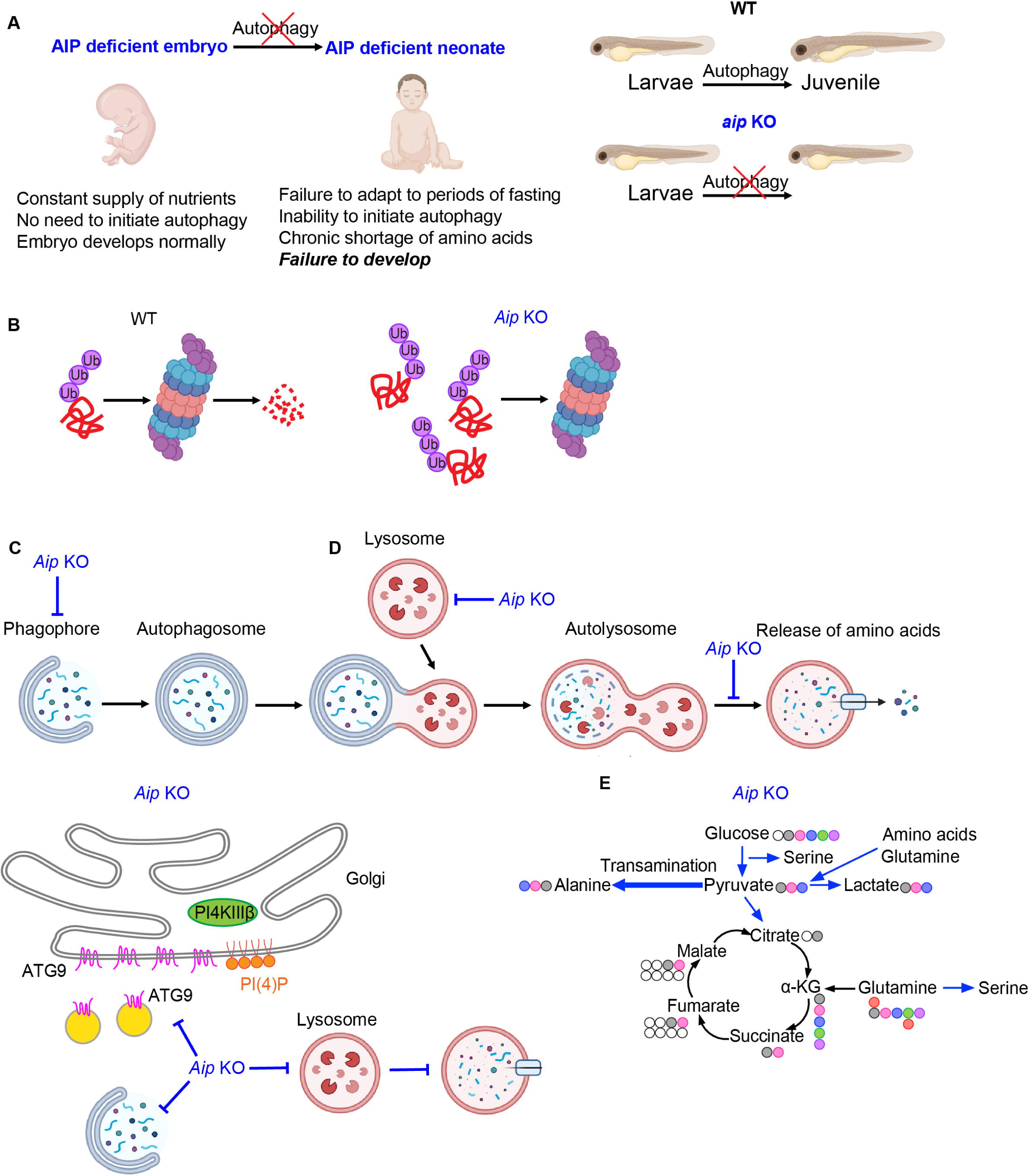
Figure S11

## Notes

### Competing Interest Statement

The authors have declared no competing interest.

