## Supplementary Material for "Biallelic Loss of Molecular Chaperone Molecule AIP Results in a Novel Severe Multisystem Disease Defined by Defective Proteostasis"

### Materials and Methods

#### Identification of homozygous Variants in AIP

Whole exome sequencing was performed for Family A using SeqCap EZ Exome Probes v3.0 (Roche) on an Illumina HiSeq2500 system with an average depth of coverage of 125×. Given the consanguineous pedigree and the similar phenotype in both probands, an autosomal recessive condition was suspected. Genetic analysis focused on homozygous variants in exonic regions or splice sites with an ultra-rare allele frequency in the population (allele frequency <0.001%) (using gnomAD version 4.1.0) that are shared between the probands. Whole genome sequencing analysis was performed in Family B, C and D due to the lack of diagnosis for their severe abnormalities using DeCode genetics (https://www.decode.com).

#### Cell Culture

MEFs (ATCC) and human dermal fibroblasts (from biopsy of the lower arm) were grown in high glucose (25mM) DMEM supplemented with 10% FBS and penicillin and streptomycin (ThermoFisherScientific) unless otherwise stated for experimental purposes. Cells were regularly checked to ensure that they were mycoplasma negative. To induce autophagy, cells were washed 3 × in PBS and Earle’s Balanced Salt Solution (EBSS) was added to cells for 2 hours unless otherwise stated.

#### Protein half-life analysis

In vitro protein half-life analysis was performed as previously described (Hernández-Ramírez *et al.*, 2016). *AIP* variants were created using site-directed mutagenesis on MYC-tagged pcDNA3.0 plasmid vector containing the human *AIP* sequence (NM_003977.4), primers available on request. The wild-type and variant AIP vectors were transfected into HEK293 cells using lipofectamine 3000 (ThermoFisherScientific). 24 hours later, transfected HEK cells were treated with 50µg/mL cycloheximide (CHX) for 0, 6, 12 and 24 hours and Western blot analysis performed.

#### Cell Viability

Cells (40,000/well) were plated in triplicate in a 96-well plate. The viability of cells the day after plating (time 0) and at 24, 48 and 72 hours was measured using a cell viability kit (CellTiter-Glo, Promega). Viability was calculated as a percentage compared to day 0.

***qPCR***

RNA was extracted from cells following treatment using TRIzol (Invitrogen). cDNA was made using High-Capacity cDNA reverse transcription kit (Applied biosystems) and used with Brilliant III UltraFast SYBR green qPCR master mix (Agilent) on the Agilent AriaMx PCR system. The PCR cycling program implemented was 1 cycle of denaturation for 3 minutes at 95°C, proceeded by 40 cycles of denaturation for 20 seconds at 95°C, and annealing for 20 seconds at 60°C.

#### Measuring Mitosox/mitotracker expression

MEFs (50,000/well) were seeded onto a 24-well plate on the day prior to the experiment. On the day of the experiment, MEFs were washed × 2 in PBS before being incubated with Mitosox/Mitotracker Green/Mitotracker Red according to the manufactures instructions. Samples were acquired on an LSR Fortessa (BD Biosciences) and the data analyzed using FlowJo software version 9.3.1 (Tree Star, Inc, Ashland, USA).

***Measuring glucose uptake using 2-NBDG***

### MEFs (50,000/well) were seeded onto a 24-well plate on the day prior to the experiment. On the day of the experiment, MEFs were washed × 2 in PBS before being incubated with glucose free DMEM supplemented with 10% dialyzed FBS for 1 hour at 37°C. Media was removed and DMEM with 100µM 2-NBDG (2-(*N*-(7-Nitrobenz-2-oxa-1,3-diazol-4-yl)Amino)-2-Deoxyglucose) added to MEFs and incubated for 30 minutes at 37°C. Media was then removed, and the MEFs were washed with PBS before adding trypsin to harvest the cells before analyzing the 2-NBDG uptake by flow cytometry.

#### Measuring NAD/NADH ratio

This was done as described in (Chung *et al.*, 2021).

#### Measuring mitochondrial membrane potential

MEFs (10,000/well) were seeded onto glass-bottom 24-well plates, 2-3 days before imaging. Media was replaced with phenol red free DMEM supplemented with 1mM glucose, 1mM glutamine, 10mM HEPES, adjusted to pH 7.4 and incubated with 25nM tetramethylrhodamine methyl ester (TMRE) for 30 minutes at 37 °C. Cells were imaged with an LSM 880 (Carl Zeiss) confocal microscope using Fluorescent 63×/1.40 oil immersion objective lens at 37°C. TMRE was excited with a 561nm Argon laser with an output power of 0.2mW as described in (Chung *et al.*, 2021).

#### Measuring pH using pHrodo

MEFs (100,000/well) were seeded onto 35mm MatTek dishes the day prior to the experiment. pH Calibration was established using pHrodo Red AM prepared in cell loading solution (ThermoFisherScientific) supplemented with 20µM nigericin and 20µM valinomycin adjusted to different pH values (4.5, 5.5, 6.5, 7.5). On the day of the experiment, MEFs were washed × 2 in live cell imaging solution (ThermoFisherScientific) and incubated with the pH adjusted pHrodo solutions for 5 minutes at 37°C before live cell imaging to produce a calibration curve. Cells were incubated with dextran pHrodo for 15 minutes at 37°C before live cell imaging.

#### DQ-OVA/Lysotracker

MEFs (50,000/well) were seeded onto coverslips the day prior to the experiment. DQ-OVA (ThermoFisherScientific) was prepared according to the manufactures instructions and incubated with cells for 15 minutes at 37°C before removing and washing the cells × 2 in PBS and adding fresh media for 30 minutes before adding lysotracker for 15 minutes.

***Amino acid, glutamine, glucose withdrawal***

MEFs were washed × 3 in PBS and media without amino acids (EBSS) added. For amino acid stimulation, cells were starved of amino acids for 3 hours and glutamine (4mM final concentration), leucine (0.8mM final concentration) added in the presence of 3% dialyzed FBS for the indicated times.

#### Metabolomic analysis

MEFs (10^6^/well) were seeded onto 6-well plates (5 × replicates and 1 for cell counting) and the next day MEFs were washed × 3 in glucose free media and 2mL glucose free media supplemented with 10% dialyzed FBS with 25mM uniformly labelled glucose (U-^13^C_6_) or 4mM uniformly labelled carbon glutamine (U-^13^C_5_) or nitrogen labelled glutamine (^15^N_2_) was added to the MEFs and incubated overnight (16 hours). MEFs were washed × 3 in ice cold PBS prepared with ultrapure water. MEFs were counted using a Denovix cell counter and 1mL of extraction buffer added/10^6^ cells. Extraction buffer (50% methanol, 30% acetonitrile, 20% ultrapure H_2_O, 50ng/mL HEPES made with LC-MS grade reagents). Cells were removed on ice and lysates added to Eppendorf tubes and agitated for 15 minutes at 4°C and then incubated for 1 hour at -20°C. Lysates were centrifuged at 13,000rpm for 30 minutes at 4°C and the supernatant collected and added to autosampler tubes and stored at -80°C. LC-MS analysis was performed using a Q Exactive Hybrid Quadrupole-Orbitrap mass spectrometer coupled to a Dionex U3000 UHPLC system (ThermoFisherScientific). The liquid chromatography system was fitted with a Sequant ZIC-pHILIC column (150mm × 2.1mm) and guard column (20mm × 2.1mm) and temperature maintained at 45°C. The mobile phase was composed of 20mM ammonium carbonate and 0.1% ammonium hydroxide in water (solvent A) and acetonitrile (solvent B). The flow rate was set at 200l/minute. The mass spectrometer was operated in full MS and polarity switching mode. The acquired spectra were analyzed using Xcalibur Qual Browser and Xcalibur Quan Browser software (ThermoFisherScientific). For metabolomic analysis using zebrafish, we used the protocol derived from (Naser *et al.*, 2021). 50 fish (6 dpf) were placed in water and incubated for 24 hours in L-glucose U-^13^C_6_ [10mM], L-glutamine U-^13^C_5_ [5mM] and L-glutamine ^15^N_2_ [5mM]. Fish were anaesthetized and snap frozen and placed in Precellys homogenizer tubes with 500μl of extraction buffer and samples processed as described.

***Proteomics***

Tandem-mass-tagging (TMT) LC-MS proteomics was performed as described (Li *et al.*, 2020) using the Kings College, London proteomics facility <https://www.kcl.ac.uk/research/facilities/proteomics-facility>. Each aliquoted sample was digested by following a routine in-solution digestion protocol prior to subsequent analysis by mass spectrometry, including in-solution reduction, alkylation and digestion with trypsin. Cysteine residues were reduced with dithiothreitol and derivatized by treatment with iodoacetamide to form stable carbamidomethyl derivatives. The digestion was carried out with a mixture of trypsin and LysC overnight at room temperature after initial incubation at 37^o^C for 2 hours. The digested peptides are cleaned up with a spin column of PR-C18 resins. The cleaned peptides were resuspended in 100mM Triethylammonium bicarbonate (TEAB) for TMTpro tag labelling. TMTpro reagents (Thermo-Fisher; Lot #: XB341490/YB370079) were added to the peptides along with acetonitrile to achieve a final acetonitrile concentration of approximately 30% (v/v) in a total volume of 100µL. Following incubation at room temperature for 1 hour, the reaction was quenched with hydroxylamine to a final concentration of 0.3% (v/v), after labelling efficiency check, Finally, 16plex tags were combined as one TMTpro set and dried using a vacuum centrifuge (Savant SPD131DDA SpeedVac, Thermo Scientific) for the next step. In order to explore a depth of proteome, a fractionation of TMTpro labelled peptide mix prior to LCMS analysis was performed using Pierce High pH Reversed-Phase Peptide Fractionation Kit. In brief, the TMT labelled peptide mixture was fractionated into a total of 8 fractions. Peptide fractions are dried using a vacuum centrifuge and resuspended in 2 % acetonitrile, 0.05 % trifluoroacetic acid (TFA) (aq) and ready for LCMS analysis. At this point, each TMTpro set generated eight fractions labelled from F1 to F8. Chromatographic separation was performed using an Ultimate 3000 NanoLC system (ThermoFisherScientific). Peptides were resolved by reversed phase chromatography on a 75µm*50cm C18 column using a three-step gradient of water in 0.1% formic acid (A) and 80% acetonitrile in 0.1% formic acid (B). The gradient was delivered to elute the peptides at a flow rate of 250nL/minute over 100 minutes. The eluate was ionized by electrospray ionization using an Orbitrap Fusion Lumos (ThermoFisherScientific, UK) operating under Xcalibur v4.4. The instrument was programmed to acquire MS data using “Synchronous Precursor Selection with Multi-notch MS3” method (SPS MS3) by defining a 3s cycle time among a full MS scan, MS/MS fragmentation and MS3 fragmentation. Data was acquired using one full-scan MS spectrum at a resolution of 120,000 at 200m/z with 100% normalized AGC target and a scan range of 400~1500m/z and maximum injection time 100 ms. The MS/MS fragmentation was conducted using collision-induced dissociation (CID) and quadrupole ion trap analyzer. Parameters were set up as 100% normalized AGC target, NCE (normalized collision energy) 35, q-value 0.25, isolation window 1.2 Th and maximum injection time 50 ms. The MS3 scan was analyzed using higher-energy collision dissociation (HCD) and Orbitrap analyzer with a synchronous precursor selection. Parameters include number of SPS 10, NCE 65, 400% normalized AGC target, maximum injection time 105 ms, resolution 50,000 at 200 Th, isolation window 1.6. 2-6 charged states are defined within this method. After LCMS acquisition, 8 raw files from one TMTpro set were generated and used for further analysis. Raw mass spectrometry data are processed into peak list files using Proteome Discoverer (ThermoScientific; v2.5) (PD 2.5). Processed data was then searched using Sequest search engine embedded in PD 2.5, against the current version of the reviewed Swissprot mouse databases downloaded from Uniprot (<http://www.uniprot.org/uniprot/>).

#### Electron microscopy

Cells were fixed 2.5% glutaraldehyde buffered with 100mM sodium cacodylate (pH7.2) for 24 hours at room temperature and stored for up to a week at 4ºC. Sample processing and imaging were performed by the Histopathology Department (Great Ormond Street Hospital, London). Briefly, after secondary fixation in 1% osmium tetroxide (agar scientific) for 2 hours, samples were dehydrated in graded ethanols, transferred to propylene oxide then infiltrated and embedded in Agar 100 epoxy resin. Polymerization was carried out at 60°C for 48 hours. Ultrathin sections (90nm) were cut using a Diatome diamond knife on a Leica EM UC7 ultramicrotome. Sections were collected on copper grids and stained with alcoholic uranyl acetate and lead citrate. Sections were examined with a JEOL 1400 TEM and digital images recorded using an AMT XR80 digital camera. Approximately 50 images of WT and *Aip* KO MEFs and ~50 images of HC and PDSF taken. Autophagosomes and lysosomes were determined using the method previously described (Huotari and Helenius, 2011).

#### Lentivirus transduction

AIP was cloned into pLOC RFP IRES GFP (Dharmacon). This plasmid was transfected into HEK293T cells with packaging plasmids using a second-generation lentiviral system. Lentivirus was collected on day 4 and day 5 post transduction and concentrated using ultrafiltration columns (Sartorius). *Aip* KO MEFs were transduced with virus and GFP positive cells were isolated by cell sorting.

#### Western blotting

Whole cell lysates were prepared following treatment. Cells were washed 3 × with PBS and lysates generated using Phospho-Safe extraction buffer (Millipore) with complete protease inhibitors (Roche) or Ripa lysis buffer and incubated for 1 hour on a rotating wheel at 4°C. Samples were centrifuged at 17,000rpm for 30 minutes at 4°C, and supernatants quantitated using Bradford protein assay kit (Biorad). To examine ubiquitylated proteins, the lysis buffer was mixed with DUB inhibitors N-Ethylmaleimide (NEM) 10mM and PR-619 50µM. 6 × Laemmli SDS loading buffer was added and samples boiled at 95°C for 10 minutes, then run on a NuPAGE 4-12% Bis Tris gel in MES SDS running buffer (Invitrogen). Proteins were transferred using semi-dry transfer either to nitrocellulose blotting membrane 0.45µm (Amersham Protran) or PVDF membrane 0.45µm (Immobilon-FL) for total and phospho-proteins respectively. Membranes were blocked in either 5% milk powder or 5% BSA for total and phospho-proteins. Antibodies (Resources) were used at a dilution of 1:1,000 for immunoblotting and incubated overnight at 4°C. The following day membranes were incubated with the corresponding IRDye secondary antibodies (LICOR) for 1 hour. Proteins were detected using the Odyssey Infrared Imaging System. Band densitometry was performed using ImageJ.

#### Confocal microscopy

Cells (50,000) were added to coverslips and the next day used for experiments. Cells were fixed with 4% PFA for 15 minutes and washed 3 times using PBS. Cells were permeabilized with 0.1% Triton X-100, 20µM digitonin, 0.5% saponin or methanol depending upon the target of interest. Coverslips were blocked with 5% BSA for 45 minutes before incubating the cells with primary antibodies in 1% BSA for 45 minutes. Coverslips were washed × 3 and secondary antibodies in 1% BSA added and incubated for 45 minutes. Cells were counterstained with DAPI-mounting solution and placed on slides. Images were taken using a LSM880 (Zeiss) with a 63× objective. After acquisition, images were processed using ZEN software provided by Zeiss. Images were quantified using Image-J Software with the mean of 3-6 separate images (30-60 cells) used with the same threshold across all samples.

#### Staining phosphoinositides

PI3P and PI4P were stained using a protocol previously described (Judith *et al.*, 2019). Cells were permeabilized for cells 5 minutes with 20µM digitonin in buffer A (20mM Pipes, pH 6.8, 137mM NaCl, and 2.7mM KCl). Cells were then washed × 3 in buffer A and then blocked for 45 minutes in buffer A containing 5% BSA. PI3P (GST-2×FYVE) (10µg) was added in buffer A for 60 minutes at room temperature. Cells were washed × 3 with buffer A, anti-GST 488 in buffer A added to cells and incubated for 45-60 minutes. Cells were post-fixed for 5 minutes with 2% PFA, washed × 3 with PBS containing 50mM NH_4_Cl and once with MilliQ water and mounted.

#### RNA sequencing

The primary analysis pipeline for RNAseq consists of FastQC version 0.11.9 to look at the quality control within the reads data. HISAT2 version 2.2.1 (Kim, Langmead and Salzberg, 2015) is used for read alignment. Sam Tools version 1.10 was used to convert SAM files to sorted and indexed bam files and Picard Metrics version 2.26.6 for further quality control. Quantification of the RNAseq data as counts per gene was completed using subread feature Counts version 2.01 (Liao, Smyth and Shi, 2014). R studio (R version 4.2.2) was used for differential gene expression analysis using DESeq2 version 1.38.3 (Love, Huber and Anders, 2014).

***Generation of aip loss of function zebrafish***

To generate *aip* knock-out zebrafish, a crRNA was designed targeting *aip* exon 2 using CHOPCHOP (https://chopchop.cbu.uib.no), where the target site was included in the recognition site of BstX1 restriction enzyme [**CCA**GGACGATGGGAGGTCACAGT (PAM sequence in bold, restriction site underlined)]. 1nL of a solution containing 62.5 ng/mL crRNA (Sigma-Aldrich/Merck), 62.5 ng/mL tracrRNA (TRACRRNA05N, Sigma-Aldrich/Merck), and 5 mM Cas9 (M0386M, NEB Ltd), was injected in one-cell stage zebrafish embryos (wild-type, TU). Around 100 embryos were injected and the knockout efficiency was assessed by polymerase chain reaction (PCR) from genomic DNA (*aip*_Forward, 5’-TCATTACCGCACTAGCCTGT- 3’; *aip*_Reverse, 5’- TGAATTCAGCTATTTCCCCCTCTT -3’), followed by BstX1 restriction enzyme digestion and 3% gel electrophoresis. The higher percentage of non-digested PCR products indicated higher efficiency and lower mosaic rate. Around 50 injected fish were raised to adulthood and crossed with wild-type TU fish to identify founders. To genotype, either the whole larva was collected individually in a 96-well PCR plate, or adult fish were anaesthetized with MS-222 at 168mg/L (E10521, NEB Ltd) and a caudal fin clipping was collected for genomic DNA extraction and amplified by PCR primers mentioned above. The amplicons were sequenced at Source Bioscience, UK. The molecular and morphological experiments were conducted among the third generation (F3) including *aip* wild-type, heterozygous and homozygous fish.

***Zebrafish maintenance***

Zebrafish were housed in a recirculating system (Tecniplast, UK) on a 14-hour/10-hour light/dark cycle and a constant temperature of 28ºC. Fish were fed twice daily with ZM-400 fry food (Zebrafish Management Ltd., UK) in the morning, and brine shrimp in the afternoon. Breeding was set up in the evening, either in sloping breeding tanks (Tecniplast, UK) or in tanks equipped with a container with marbles to isolate eggs from progenitors. For experiments where the developmental stage of larvae was important, we placed barriers between the fish to keep them isolated in the breeding tank. The following morning, barriers were removed to allow spawning. Eggs were collected in 90mm Petri dishes the following morning, sorted fertile from infertile, and then incubated at 28ºC (max 50 eggs/dish). Dishes were checked daily to ensure consistent developmental stage across groups. If reared, larvae were moved to the recirculating system at 5 days post fertilization (dpf). All procedures were carried out under license in accordance with the Animals (Scientific Procedures) Act, 1986 and under guidance from the Local Animal Welfare and Ethical Review Board at Queen Mary University of London.

***Morphological analysis***

From 5 dpf all larvae were assessed twice daily for effects on morphology or behavior. Any larvae showing altered morphology such as yolk sac edema, body curvature, altered swimming or lack of movement were killed in accordance with UK animal legislation. For size measurements, zebrafish larvae were anaesthetized using MS-222 at 168mg/L and mounted in 3% methyl cellulose (M7027, Sigma-Aldrich/Merck) in E3 embryo medium. Images were taken by a Leica S9i stereo microscope with an integrated 10 M pixels network camera. The body length was measured using the free graphical analysis software ImageJ (National Institutes of Health, Bethesda, Maryland, US). Following imaging, larvae were washed to remove methyl cellulose and transferred individually to single wells of a multi-well plate or single tanks for genotyping.

Zebrafish larvae were anaesthetized using MS-222 at 168mg/L and mounted in 3% methyl cellulose (M7027, Sigma-Aldrich/Merck, UK) in E3 embryo medium. Images were taken by a Leica S9i stereo microscope with an integrated 10 M pixels network camera. The body length was measured using the free graphical analysis software ImageJ. Following imaging, larvae were washed to remove methyl cellulose and transferred individually to single wells of a multi-well plate for genotyping.

***Genotyping of larval fish***

For experiments where no further analysis was necessary after genotyping (morphological analysis and survival experiments), genomic DNA was extracted from the whole larvae with 100μL of 50mM sodium hydroxide (NaOH, 655104, Sigma-Aldrich/Merck) before incubation at 95°C for 30 minutes, followed by the addition of 10μL of 1mM Tris-HCl (10812846001, Sigma-Aldrich/Merck). For immunohistochemical analysis, individual larvae were fixed with 4% paraformaldehyde (PFA, 158127, Sigma-Aldrich/Merck), and then only tail sections were collected and incubated in 25μL of 50mM NaOH at 95°C for 20 min, followed by the addition of 2.5μL of 1mM Tris-HCl. The genotyping PCR primers and procedure were as described above.

***Immunofluorescence***

Fixed and genotyped larvae were washed briefly with phosphate buffered saline (PBS, P4417, Sigma-Aldrich/Merck). Samples were cryoprotected in 15% and 30% sucrose overnight at 4°C before being embedded in optimal cutting temperature compound (OCT, AGR1180, Agar Scientific Ltd, UK) in a cold bath with dry ice. Sectioning of blocks at 12μm thickness was performed using a cryostat (CM3050S, Leica Microsystems, UK) and mounted on SuperFrost Plus microscope slides (ThermofisherScientific). Primary antibodies including Anti-Ubiquitin 1:250 (ab134953, Abcam) and anti-LC3B 1:200 (ab192890, Abcam) and goat anti-rabbit secondary antibody Alexa Fluor™ 546 (11035, Invitrogen) were used for immunofluorescence following standard protocol (Ferguson and Shive, 2019). All slides were counterstained with DAPI (1μg/mL) for 10 minutes.

***Zebrafish metabolomic analysis***

To avoid the need to genotype all larvae, as homozygous larvae could be identified based on morphology, metabolomic analysis was performed on groups of homozygous mutants identified from F3 heterozygotes in-cross while wild-type controls were obtained from F3 wild-type siblings in-cross. Groups of 50 fish were raised to 5 dpf in E3 medium with 10mM labelled glucose (U-^13^C_6_) or 5mM carbon labelled glutamine (U-^13^C_5_) overnight at 28°C for 24 hours. The next morning, fish were briefly washed with cold PBS and extracted as described for MEFs.

***Assessing autophagy in larval zebrafish***

The induction of autophagy was assessed following acute dietary restriction. Zebrafish larvae were reared under normal conditions until 5 dpf. At 5 dpf WT and homozygous mutant larvae were separated into 2 groups for normal and restricted feeding. The normal group was fed normally twice a day with paramecium from 5-7 dpf while the restricted group were fed normally at 5 dpf but received no further paramecium before being harvested for protein analysis.

***Dietary restriction conditions***

Larvae were randomly divided into two groups at 5 dpf and each group included six biological replicates with 25 embryos each. The control group was fed with paramecium twice a day. The restricted group was fed normally on day 5 but received no further paramecium. All other rearing conditions were as normal. Larvae were checked twice a day for signs of harm, such as yolk sac edema, body curvature, and lack of movement even upon stimulus. Any larvae showing these symptoms were humanely euthanized and recorded as dead. After euthanasia, the entire fish was collected for genomic DNA extraction and genotyping. Larvae were reared until 7 dpf and number of larvae removed each day recorded.

***Zebrafish Western blotting***

Zebrafish larvae were reared under normal fed conditions until 5 dpf. They were fed in the morning of the 6^th^ dpf and then harvested for Western blot analysis in the morning of the 7th dpf. Then larval fish were put in 200uL cold RIPA lysis buffer (20-188, Sigma-Aldrich/Merck) with 1mM PMSF. Samples were then homogenized using a bead mill homogenizer and incubated on a shaker for 30 minutes at 4°C before centrifuging at 13,000rpm, 4°C for 15 minutes. 150μL supernatant in each sample was transferred to a clean 1.5mL tube and performed Bradford assay (B6916, Sigma-Aldrich/Merck) to determine concentration.

After boiling for 10 minutes, 15μg protein was loaded in each lane of 12% Mini-PROTEAN® TGX™ Precast Gels (4561043, Bio-Rad) and separated in 1× Tris-Glycine Buffer (T4909, Sigma-Aldrich/Merck), followed by transferring protein to membrane using Trans-Blot® Turbo™ Transfer System and Trans-Blot® Turbo™ Mini PVDF Transfer Packs (1704156EDU, Bio-Rad). After transferring, the blots were blocked with 2.5% skimmed milk in PBST, followed by incubation first with the Anti-LC3B antibody 1:1000 (ab192890, Abcam) and Anti-beta actin antibody 1:1000 (ab8227, Abcam), and then probed with Goat Anti-Rabbit IgG H&L (HRP) (ab205718, Abcam) and visualized with SuperSignal™ West Pico PLUS (34590, ThermoFisherScientific). NIH software Image J was applied for blot scanning and protein area quantifications.

***Proteasome activity***

Proteasome activity was analyzed using Proteasome Activity Assay Kit (ab107921, Abcam, UK). Groups of wild-type and homozygous larvae were generated as described above for metabolomic analysis. 10 fish from each group were homogenized in 200µl 0.5% NP-40 in distilled water by pipetting up and down and then centrifuged for 15 minutes at 13,500 rpm, 4°C. According to the manufacturer's instruction, samples were incubated with an AMC-tagged peptide substrate for 30 minutes at 37° C to produce fluorophore. Finally, the fluorescence was measured by using FLUOstar Omega Microplate reader (BMG LABTECH) at Ex/Em = 350/440 nm in kinetics mode at 37°C every ten minutes in 60 minutes.

**LysoTracker staining**

LysoTracker staining was performed as previously reported (He and Klionsky, 2010). Larvae were incubated in 5µM LysoTracker Deep Red (L12492,ThermoFisherScientific) in E3 medium for 2 hours with gentle shaking and washed ×3 with E3 medium. Live images were acquired with Leica fluorescence stereomicroscope MZFLIII (Leica).

| Resources |  |  |
| --- | --- | --- |
| **Antibodies** | **Company** | **Catalogue #** |
| Aldolase A | Cell Signaling Technology (CST) | 8060 |
| ULK1 | CST | 8054 |
| Phospho-ULK1 (S555) | CST | 6869 |
| AMPK1α | Proteintech | 66536-1 |
| Phospho-AMPK1 (Thr172) | CST | 2535 |
| p62 | CST | 5114 |
| Phospho-p70 S6K (Thr389) | CST | 975965 |
| p70 S6K | CST | 2708 |
| RAB7 | Abcam | ab126712 |
| Ubiquitin | Abcam | ab134953 |
| Linkage specific K48 Ubiquitin | Abcam | ab140601 |
| LAMP1 | BD Biosciences | 553792 |
| TFEB | ProteinTech | 13372-1AP |
| TFE3 | ProteinTech | 14480-1-AP |
| LC3B | Abcam | ab192890 |
| LC3 | Abcam | ab192890 |
| Wipi2 | Invitrogen | PA5-54098 |
| GM130 | BD Biosciences | 610823 |
| ATG9a | Invitrogen | MA1-149 |
| RAB7 | Abcam | Ab126712 |
| PI(4)P | Echelon Bioscience | Z-P004 |
| PI4K3β | BD Biosciences | AB_399297 |
| NPC2 | Invitrogen | PA5-51463 |
| GAA | Abcam | ab314448 |
| Total Oxphos antibody cocktail | Abcam | ab110413 |
| AIP (Xap2) | Santa Cruz | Sc-59730 |
| Plasmids |  |  |
| GST-2×FYVE | Gift from Bart Vanhaesebroeck, UCL |  |
| Cell lines | **Company** | **Catalogue #** |
| *Aip* knockout mouse embryonic fibroblasts | ATCC | CRL-3288 |
| Wild-type mouse embryonic fibroblasts | ATCC | CRL-3287 |
| Healthy control dermal fibroblasts |  |  |
| Patient derived dermal fibroblasts |  |  |
| Culture reagents |  |  |
| DMEM high glucose | Sigma | D6429 |
| Earle’s Balanced Salt Solution (EBSS) | ThermoFisherScientific | 24010043 |
| Penicillin-Streptomycin | ThermoFisherScientific | 15140130 |
| Glutamine free media | ThermoFisherScientific | 11960044 |
| Glucose free media | ThermoFisherScientific | 11966025 |
| FBS | ThermoFisherScientific | 26140079 |
| Dialyzed FBS | ThermoFisherScientific | 26400044 |
| Leucine | ThermoFisherScientific | A12311.22 |
| L-Glutamine | ThermoFisherScientific | A2916801 |
| NEAA | ThermoFisherScientific | 10370047 |
| Reagents | **Company** | **Catalogue #** |
| EGCG | Tocris | 4524 |
| L-DON | Tocris | 6809 |
| AOA | Cayman Chemicals | 28298 |
| BPTES | Tocris | 5301/10 |
| Metformin | Tocris | 2864 |
| Bafilomycin | Tocris | 1334 |
| Rapamycin | Tocris | 1292 |
| 2-DG | Sigma | D6134 |
| Glucose | Sigma | G7021 |
| Oligomycin | Sigma | 75531 |
| Bovine Serum Albumin Alexa Fluor 488 | ThermoFisherScientific | A13100 |
| Dextran Alexa Fluor 488 | ThermoFisherScientific | D22910 |
| Mitotracker Red FM | ThermoFisherScientific | M22425 |
| Mitotracker Green FM | ThermoFisherScientific | M7514 |
| MitoSox | ThermoFisherScientific | M36008 |
| 2-NBDG | ThermoFisherScientific | N13195 |
| LysoTracker Red DND99 | ThermoFisherScientific | L7528 |
| DQ-OVA | ThermoFisherScientific | 10023372 |
| Dextran pH Rodo | ThermoFisherScientific | P10361 |
| Ammonia assay kit | Abcam | ab83360 |
| Proteasome assay kit | Abcam | ab107921 |
| Cell viability kit (CellTiter-Glo) | Promega | G7571 |
| Intracellular pH calibration kit | ThermoFisherScientific | P35379 |
| Labelled Metabolites | **Company** | **Catalogue #** |
| D-Glucose U-13C6 | Cambridge Isotope Laboratories Inc | CLM-1396 |
| L-Glutamine U-15N2 | Cambridge Isotope Laboratories Inc | NLM-1328 |
| L-Glutamine U-13C5 | Cambridge Isotope Laboratories Inc | CLM-1822-H |
| **Stock images** | https://biorender.com |  |

#### Mouse Primers used

| **Gene** | **Forward** | **Reverse** |
| --- | --- | --- |
| *Pik3ca* | TCTTCCCCAGAACTGCCAAAG | AGCTGCTCAGAGGACAACAAC |
| *Pik3c2a* | ATGACGAGGTGGCAGCTTTT | TGGAGACCTTCACACTGGCA |
| *Pik3cg* | CAGCAGGTCCATGTGATTGAGA | CCGGCTTTTAGTCCAGGATCAT |
| *Pik3r4* | GCCACCGAGTTAGAGTACATGAG | GCAGCTGAGAGAGATCAAACAAAG |
| *Rab5* | GACCTTCCACAAGACAGGCC | CCACGGGTCTGCATTCTTTC |
| *Rab7* | GCCACAATAGGAGCGGACTT | GGGCAGTCACATCAAACACC |
| *Atg2b* | GAGCCACTCTACAGCACAGG | AGGGGCCTGTAATCAAGTGC |
| *Prkaa2* | ATGGTTGTCCATAGGGACCTG | GCGATCCACAGCTAGTTCGTAG |
| *Them55B* | CCATGATTACCTGTCGCGTC | GACGCGACAGGTAATCATGG |
| *Becn1* | GGAGGGGTCTAAGGCGTC | GTTAGCCTCTTCCTCCTGGG |
| *Ctsl1* | GAGTGGCACCAGTGGAAGTC | CCATTCACCACCTGCCTGAA |
| *Atp6v1h* | AGATGGACATTCGAGGTGCTG | CTTGCTTGTCCTCGGAACTTC |
| *Rab7a* | GGGGACTCTGGTGTTGGAAAG | CTGGAACCGTTCTTGACCGG |
| *Lamp1* | GCATACCGGTGTGTCAGTG | GATACAGTGGGGTTTGTGGG |
| *Sqstm1* | GGCACAGAAGACAAGAG | CATTTGCTCTGTCAGAGAC |
| *Vps11* | GACCACCCACTTGTTTGTCG | CACACTCATCCCCAGCAACA |
| *Uvrag* | CAGCTTCTGGATACCTACTTCACTC | TCCACCCCAAATCTTCACCAC |
| *Atg9b* | CCCTTGCTTCATGTACCCG | CAGCGAAGGAGGAAGGTTG |
| *Wipi1* | CTGCACATCCCTAGCGATTG | GCCGAGGTTTTGTGTGACTG |
| *Ctsf* | GCTGACACTGCTCTCGACTG | CTCCTAGCACGGCCCTTGTC |
| *Atp6v0e1* | TCGTGATGAGCGTGTTCTGG | CAGAGGATTGAGCTGTGCCA |
| *Mcoln1* | GTCGGTGTCATTCGCTACCTG | GAACGATCCAGCCACAGAAGC |
| *Tfeb* | AGCAGGAGGAGCAGCGAGAG | GGAAGTGGACAGGGGTGTTG |
| *Wipi2* | ACACGTCCCTAGCTGTTGGT | GCTTTGAGGCTGACAATGGC |
| *Ctsd* | GCCCCATCACCAAGTACTCC | CCCACAGGTTAGAGGAGCCA |
| *Atg9a* | CGAGTTTGCCCAGCTCTTCC | TATGATCTCCAGGGCCCGAG |
| *Lamp2* | TCCTAGGAGCCGTTCAGTCC | GCACTATTCCGGTCATCCCC |
| *Atg13* | CGACTCTCCAGGAAGCAAGG | CAGGAAAGAGTGAGGGTGCC |
| *Atg14* | CTGTTGAGCGGTGTCCTCTG | GCCGCTTTCCTTCCATAGCT |
| *Atg16l1* | CAGAGGCAGGCGTTCGAGGA | CCACGCACCATCATGTCCAG |
| *Atg3* | GAAGTGGCCGAGTACCTGAC | GCCTTCACTTTCAATTCTTCCCC |
| *Atg7* | TTGACGTTGGAGTTCAGTGCT | AAGCCTCAAGTGTGTTGGTGT |
| *Vps34* | CGAGTGGCTGAAACTTCCGG | CAAGTCATGCATTCCTTGGCG |

#### Human Primers used

| **Gene** | **Forward** | **Reverse** |
| --- | --- | --- |
| ATG9a | CCAGAACTACATGGTGGCACT | GTCCCCAGAAGAGGATCAGC |
| LAMP2 | AATGCCACTTGCCTTTATGC | GGTCCGAACTGCACTGCTAT |
| ATG13 | GTCATTGCTGCTGAAGTCCCT | GCCACTCAGCTGAACTTCTCC |
| VPS34 | TTCCTGTGGCTGAAGTTCTTGAT | AAGTGTCCATGACCTCAGCAC |
| TFEB | GGAACAAGTTTGCTGCCCAC | GACATCATCCAACTCCCTCTCAG |
| ATPVO6E1 | GGCTTCGTCGGCTTCTTGGT | TTGCGGTCCAAAGAGAGGGT |
| ATG9b | GTAGGCCCTTTGATCCCTGA | TGTCCAGGTTCTGGATGTGA |
| MCOLN1 | ATGACACCTTCGCAGCCTAC | GGTAGTACCGCTGGCAGAGA |
| SQSTM1 | GGAACCCGCTACAAGTGCAG | TGTCCGTGTTTCACCTTCCG |
| UVRAG | CCCTTCCCAACCTGAAAAAC | CCGTTTTCGATGTCCTTTGT |
| LAMP1 | GACACCAAGAGTGGCCCTAA | CTCATGAGCTGGACGCTGTA |
| CTSF1 | AAGAGCCTGGCAACAAGATG | CCACATTGCCTGTGACTGAG |

#### Statistical analysis

Statistics were performed using Graphpad Prism version 9.2 software. Comparisons were made with Student’s *t*-test, Mann-Whitney test, 1-way and 2-way ANOVA. Significance was taken as p<0.05 and marked on figures as *p <0.05, **p <0.01, ***p <0.001, ****p <0.0001. Error bars on figures represent standard error (SEM).

**Supplementary material**

#### Patients

Clinical parameters are detailed in **Supplementary Table S1.**

Four of the five patients required neonatal intensive unit care after birth due to extreme weight loss, with failure to thrive despite full nutritional support. All had persistent tachycardia, profuse diarrhea and mild hyponatremia, with pronounced metabolic acidosis. They had intermittent hypoglycemia (lowest value in Patient FCM1 2.86 mmol/l), as well as intermittent hypertonia, and gastrointestinal bleeding with anemia in three patients. Two patients had septal atrial defects and one a persistent ductus arteriosus and systolic murmur. It should be noted that *Aip* deficient mice have heart developmental abnormalities including sepal defects and patent ductus arteriosus (Lin *et al.*, 2007). Two patients had some dysmorphic features: FBM1 was noted to have a long and slender body, long fingers and toes, while FCM1 had a triangular face with dysplastic low-set ears, funnel chest, and later craniosynostosis with a small head circumference. All patients had a very low BMI (See Table).

During hospitalization, the main care challenges included feeding problems with recurrent sub-obstruction with subsequent ileostomy formation in one child, diarrhea, failure to thrive despite total parenteral nutrition (TPN) support, episodes of elevated blood pressure (up to 130/80mm Hg), tachycardia (up to 160/minute). One of the major challenges for all five patients was to deal with episodes of unexplained hyperthermia: axillary temperatures ranged usually between 36.5C and 39.8C with no sign of infection (**Fig S1A**). This was repeatedly investigated by microbiological assessment of various body fluids, but without positive results, and treated empirically with antibiotics; however, the episodes were not accompanied with elevated leukocytes or CRP, and screenings including bacterial and viral serology were also negative. Occasionally, a higher temperature was correlated with an elevated CRP, suggesting true short-term infection, but this was an uncommon finding. All patients showed general high levels of irritability and agitation, with episodes of monotonous loud crying, hyperesthesia and severe hyperhidrosis. They demonstrated arching with head tilted back, seemingly in pain and requiring intermittent sedation. Circulating catecholamines were normal in all 5 patients. Anemia was present in all patients: FCM1 received erythropoietin treatment. Micropolyadenia was noted in two of the 5 children. Hepatomegaly was seen in three patients (1.5cm in FCM1). Cardiac ultrasound identified left ventricular hypertrophy in children.

*Biochemistry*

Low thyroid stimulating hormone (TSH) levels with a normal free thyroxine (fT4) levels were noted in all children, with raised aldosterone, DHEAS, testosterone and prolactin levels in the one where this was tested. Serum calcium levels were high normal or high (highest calcium levels were: total calcium 3.43mmol/L [2.25-2.75] in FCM1). Hypercalciuria was present, phosphate level was high or normal (range in FCM1 was 1.98-2.56mmol/l [1.45-1.78]) and parathormone was normal or low. In some patients, vitamin D level was supplemented initially as routine but stopped when hypercalcemia was noted. Vitamin D and 1,25 Vitamin D levels were normal. Abdominal ultrasound identified nephrocalcinosis in all the children from the age of 6 weeks, and one of them had duplication of the collecting system on one of the kidneys.

*Imaging*

MRI of the brain revealed no structural abnormalities although there were potential signs of early demyelination in one case. EEG and EMG studies were both within normal limit. FBM1 and FCM1 had bilateral hydrocele. FCM1 had inguinal hernia and underwent surgery at the age of 3 months.

*Therapy*

Four patients received treatment with partial/total parenteral nutrition, repeated courses of antibiotics, erythropoietin, ACE inhibitors, calcium channel blockers, β-adrenoceptor blockers, acetylsalicylic acid for bowel irritation and thyroxine. Prednisolone was attempted to reduce calcium with biochemical success (high normal calcium and PTH rise achieved), but no clinical improvement.

*Outcomes*

FAM1 died age 8 month weighing 4285g. FAM2 passed away at age 10 months old, at that time, he weighed 4090g, with a length of 62.0cm. FBM1 is alive on TPN, has normal IGF-1 and prolactin levels, body height is between 90-97 centile with body weight 25-50 centile. He has moderate intellectual disability with relatively severe speech delay and a diagnosis of autism spectrum disorder. Body temperature elevations ceased around age 11 months. FCM1 had his last admission age 10 months, when he had height at 47 percentile (71cm), weight <0.1 percentile (6.15kg), head circumference <0.1 percentile (41cm), he was lost to follow-up after discharge at 10.5 months of age. FDM1 at the age 14 months has increased muscle tone in the limbs, high popliteal reflexes, and a Babinski sign on the left side. Mental development is delayed but he can pronounce a few words at age 20 months, and his physical condition is stable with slow weight gain, body weight -2.26 standard deviation. His body temperature has remained normal after 14 months of age.

#### Table S1 Clinical parameters of the patients

|  | **Family A**  **FAM1** | **Family A**  **FAM2** | **Family B**  **FBM1** | **Family C**  **FCM1** | **Family D**  **FDM1** |
| --- | --- | --- | --- | --- | --- |
| **Sex** | Female; XX | Male; XY | Male; XY | Male; XY | Male; XY |
| **Ethnicity** | Turkish | Turkish | Icelandic | Ossetian | Ossetian |
| **Mutation in *AIP* gene** | NM_003977.2: c.62G>A | NM_003977.2: c.62G>A | NM_003977.2: c.827>A | NM_003977.2: c.346_372 deletion | NM_003977.2: c.346_372 deletion |
| **Expected protein change** | NP_003968.2: p.Gly21Asp | NP_003968.2: p.Gly21Asp | NP_003968.2: p.Ala276Glu | NP_003968.2: p.Glu116_Val124del | NP_003968.2: p.Glu116_Val124del |
| **Current status** | Died 8 months | Died 10 months | Alive, closely followed | Lost for follow-up at 10.5 months | Alive, closely followed |
| **Birth weight, length** | 1695 grams, 44.2 cm | 2430 grams, 48.5 cm | 2426 grams, 50 cm | 3100 grams, 49 cm | 3350 grams, 52 cm |
| **Body weight** | 4285 grams at 8 months | 4090 grams at 10 months | 8820 grams at 8 months | 6710 grams at 10 months | 7800 grams at 14 months |
| **Dysmorphic features** | None | None | Long and slender body and fingers and toes, some loose skin of face, funnel chest deformity | Triangular facial shape  Funnel chest deformity  Early closure of cranial vault sutures | None |
| **Pregnancy history** | Born at 31 weeks  Premature rupture of membranes | Fetal tachycardia Polyhydramnios  Born at 36 weeks after induction | Mother had pre-eclampsia requiring early delivery at 35 weeks 4 days. From 30 weeks high blood pressure treated with labetalol and from week 35 preeclampsia, treated with magnesium for 24 hours prior to delivery | Week 37.5 abruption of placenta  Caesarean section | No complications, born at 38 weeks |
| **Genetic testing** | Standard karyotype Whole exome sequencing | Whole exome sequencing | Microarray normal, spinal muscular atrophy testing normal, panel for D vitamin metabolism genes normal, whole genome sequencing | Whole exome sequencing, CYP24A1 heterozygote VUS identified | Whole exome sequencing |
| **Cardiac** | Tachycardia, hypertension, large atrial septal defect, L>R shunt, beta blocker for tachycardia and hypertension | Tachycardia  hypertension beta blocker for tachycardia and hypertension | Tachycardia, hypertension, mild hypertrophy on echocardiography soon after birth, ongoing. On enalapril and propranolol for hypertension and tachycardia | Tachycardia, hypertension, atrial septal defect (hemodynamically insignificant), open arterial duct, left ventricular hypertrophy.  On ACE inhibitor, calcium channel blocker, beta blocker for tachycardia and hypertension | Tachycardia, normal cardiac US, normal blood pressure |
| **Pulmonary** | Normal ventilation, tachypnoea during episodes of hyperthermia | Normal ventilation, tachypnoea during episodes of hyperthermia | Normal ventilation, tachypnoea during episodes of hyperthermia | Apnea, bronchitis | Repeated bronchitis |
| **Gastrointestinal** | Episodes of diarrhea, fat in stool normal, biopsy: non-representative sample, obstruction, high calorie total parenteral nutrition (TPN), needed ileostomy age 18 days | Gastrointestinal bleeding age 7 days, episodes of diarrhea, obstruction, high calorie TPN nutrition | Chronic secretory diarrhea, on long-term TPN nutrition, gastroscopy and colonoscopy normal, gut biopsies no specific abnormality found | Episodes of dehydration, diarrhea | Hepatomegaly, splenomegaly, episodes of poorly digested stool (1-2/day) |
| **Renal** | Polyuria, aminoaciduria, hypercalciuria, normocalcemia | Polyuria, medullar nephrocalcinosis with hypercalciuria, normocalcemia | Hypercalcemia and hypercalciuria, nephrocalcinosis, kidney stones. Treated with low calcium TPN formula (70% of normal).  Bisphosphonate trial lowered blood calcium on 2 occasions of broken bones with minimal/no trauma | Enlarged kidney, hypercalcemia (max value 3.43mmol/L), hypercalciuria, nephrocalcinosis | Hyperechoic inclusions in kidneys, mild hypercalcemia (0.79 mmol/L) |
| **Kidney ultrasound** | Increased reflectivity kidney parenchyma, enlarged adrenal glands | Nephrocalcinosis | Nephrocalcinosis | Enlarged kidney Hyperechogenic kidneys  Abnormal vas deferens morphology | Hyperechogenic kidneys |
| **Neurology** | Episodes of increased tone, irritability and discomfort | Episodes of increased tone, irritability and discomfort | Episodes of increased tone, irritability and discomfort.  Has developmental delay (speech and gross motor), formally diagnosed with autism.  Large motor skills behind | Dystonia, encephalopathy, pronounced anxiety (with head tilting), delayed ability to sit, gross motor development | Episodes of increased muscle tone, rapid reflexes, positive Babinski on one side, motor developmental delay, cannot stand or sit age 14 months |
| **Temperature** | Episodes of hyperthermia >40°C | Episodes of hyperthermia >40°C | Episodes of hyperthermia, less noticeable after 11 months of age | Episodes of hyperthermia, 38-39°C | Episodes (~monthly) of hyperthermia(38-39°C) until 14 months of age. Subsequently, these episodes became less frequent |
| **Brain MRI** | Delayed myelination of the crus posteriors of the capsula interna | Repeatedly mild ventriculomegaly | Normal | Normal | Not performed |
| **Liver** | Prominent liver, normal intensity | Normal | Normal | Hepatomegaly, elevated hepatic transaminase | Hepatomegaly, transiently elevated liver enzymes |
| **Infection** | Negative | negative | Low immunoglobulins, on iv immunoglobulins from age 1.5 | Bacteriuria | Repeated bronchitis |
| **Hematology** | Iron-deficiency anemia | Anemia | Anemia, bone marrow biopsy normal, iron infusions 3x until now in his life, trying Imodium now, antihistamine trying make him sleep better | Anemia | Anemia, thrombocytosis |
| **Endocrinology** | Episodes of hyperhidrosis, normal catecholamine, VIP and chromogranin secretion  Low TSH, mild increased aldosterone,  normal DHEAS and testosterone  normal PTH  normal GH and IGF-1,  increased prolactin, intermittent hypoglycemia | Episodes of hyperhidrosis,  Normal catecholamine and cortisol secretion, low TSH, low  1,25-hydroxy VitD, normal PTH  High-normal GH and IGF-1,intermittent hypoglycemia | Episodes of hyperhidrosis, on cooling mattress and fan, body covered with  sweat droplets on skin.  Normal urine metanephrins.  Normal thyroid function, normal prolactin, FSH, LH and cortisol.  normal insulin, low PTH, Normal 1,25-hydroxy VitD  Despite normal GH and IGF-1, 2SD above mean for height since age 2 years, .  intermittent hypoglycemia | Episodes of hyperhidrosis,  normal catecholamines, low TSH,  low PTH,  height 47 percentile while weight <0.1 percentile at age 10m, episodes of hypoglycemia | Episodes of hyperhidrosis,  low TSH, normal freeT4,  normal 25-OH VitD, GH and IGF-1,  episodes of hypoglycemia |
| **Immune** | Normal | Normal | Lymphadenopathy (axilla, neck) on both sides, biopsy not concerning | Micropolyadenia (6-9 groups of the increased lymph nodes), decreased antibody level in blood | Micropolyadenia, elevated IL8 levels |
| **Metabolic** | Electrolyte disturbance because of diarrhea; amino acids plasma and urine: normal; Organic acids urine- normal; Long chain fatty acids: normal; Low free carnitine (11 µmol/L [19 – 69]) | Electrolyte disturbance because of diarrhea; Normal amin- acids in plasma, CSF and urine Normal urine organic acids  Normal sialotransferrins Normal long chain fatty acids  Normal mitochondrial enzymes | Normal plasma amino acids, lactate, acylcarnitine ammonium highest at 83, but also a value of 48 | Electrolyte disturbance because of diarrhea,  normal fatty acids.  normal amino acids | Normal amino acid and fatty acid levels,  normal lactate and ammonium |
| **Treatments received** | TPN, steroids, tacrolimus, sedatives, analgesics, beta blockers | TPN, steroids, tacrolimus, sedatives, analgesics, beta blockers | TPN, Beta blockers, ACE inhibitors | Repeated antibiotics, oral nutritional support, erythropoietin, ACE inhibitor, calcium channel blocker, beta blocker, TPN | Repeated antibiotics, oral nutritional supplements |
| **Comments** |  |  | Gets tired quickly, then lies on the floor | The family is lost from follow-up | Same ethnic background and geographical area as Family C. |

#### Table S2 Allele frequency of the three identified AIP variants in various databases

#### (this data does not include the five probands and their parents)

|  | **c.62G>A; p.Gly21Asp**  11-67483220-G-A  (GRCh38) | **c.827C>A; p.Ala276Glu**  11-67490827-C-A  (GRCh38) | **c.346_372del; p.Glu116_Val124del** |
| --- | --- | --- | --- |
| **Alleles tested in gnomAD** | 1614206 | - | - |
| **Alleles found in gnomAD** | 5 heterozygote (2 non-Finnish European, 3 far Eastern)  0 homozygote | Not identified | Not identified |
| **Allele frequency in gnomAd** | 0.000003097 | - | - |
| **Alleles tested in Iceland** |  | 319218 | - |
| **Alleles found in Iceland** | Not identified | 27 heterozygote  0 homozygote | Not identified |
| **Allele frequency in Iceland tested population** |  | 0.00008458 |  |
| **Alleles tested in Russia Genetic project** |  |  | 78985 |
| **Alleles tested in Russia Genetic project** | Not identified |  | 3 heterozygote (all 3 from the same geographical region as probands)  0 homozygote |
| **Allele frequency in Russia tested population** |  |  | 0.00003798 |

### Supplementary Figure legends

**Supplementary Figure 1**

Axillary temperature profile over a period of 4 months from patient FBM1 showing spikes in temperature coinciding with periodic fevers (**A**).

**Supplementary Figure 2**

Co-localization of ubiquitin (Ub) and LC3 under fed conditions in WT and *Aip* KO MEFs (**A**). WT and *Aip*-KO-MEFs were cultured in the absence of amino acids (EBSS), EBSS plus bafilomycin (100nM), complete media (DMEM) plus bafilomycin and the viability after 48 hours was determined (**B**). Expression of ubiquitin and Wipi2 in WT, *Aip* KO MEFs and *Aip* KO “rescue” MEFs (**C**). Induction of autophagy treating with *Aip* KO MEFs and *Aip* KO “rescue” MEFs with rapamycin (Rapa) 100nM and bafilomycin (Bafilo) 100nM for 2 hours and blotting for LC3 (**D**).

Number of PI3P^+^ (**E**), Wipi2^+^/PI4P^+^ (**F**), PI4K3β^+^/Wipi2^+^(**G**) puncta in WT and *Aip* KO MEFs under starved (2 hours EBSS). Graphs show the mean ± SEM from at least two independent experiments. Students un-paired, paired *t*-test and 2-way ANOVA were used for analysis.

**Supplementary Figure 3**

TFEB expression determined by western blotting under fed and starved (2 hours EBSS) in WT and *Aip* KO MEFs (**A**). Expression of TFE3 in fed and starved WT and *Aip*-KO-MEFs, A.U. arbitrary units (**B**). Autophagy gene expression in WT and *Aip* KO MEFs under fed conditions determined by qPCR (**C**). Relative expression of autophagy genes under fed (Time 0) and starved (EBSS) over time in WT and *Aip* KO MEFs determined by qPCR (**D**). Heat map of RNA-sequencing data showing relative expression of genes involved in autophagy (**E**). Graphs show the mean ± SEM from at least two independent experiments. Students un-paired, paired *t*-test and 2-way ANOVA were used for analysis.

**Supplementary Figure 4**

Induction of autophagy by culturing healthy control (HC) and AIPd-PDFs (FAM1, FBM1 and FDM1) in EBSS determined by immunofluorescence for Wipi2 (**A**) and Western blotting for LC3 and p62 (**B**). Induction of autophagy by treating healthy control (HC) and AIPd-PDFs with rapamycin (Rapa) 100nM and bafilomycin (Bafilo) 100nM for 2 hours (**C**). Expression of autophagy genes in HC and AIPd-PDFs (FBM1) under fed conditions determined by qPCR (**D**). Electron microscopy analysis of HC and AIPd-PDFs (FBM1) with the number of vesicles and autophagosomes determined (**E**). HC and AIPd-PDFs (FBM1) were incubated with DQ-OVA and stained with lysotracker (**F**). ­Graphs show the mean ± SEM from at least two independent experiments. Students un-paired, paired *t*-test and 2-way ANOVA were used for analysis.

**Supplementary Figure 5**

Heat map of RNA-sequencing data showing relative expression of lysosome genes under fed conditions in WT and *Aip* KO MEFs (**A**). Heat map of RNA-sequencing data showing relative expression of genes involved associated with lysosomal storage diseases. List of genes from <https://panelapp.genomicsengland.co.uk/panels/529/> (**B**).

WT and *Aip* KO MEFs were starved of amino acids (EBSS media supplemented with 3% dialyzed FBS) over time and the relative amount and puncta size of LAMP1 determined by immunofluorescence (**C**). WT and *Aip* KO MEFs were starved of amino acids (aa) for 3 hours followed by the addition of 3% albumin and the relative amount of mTORC1 activation (pS6K1 expression) over time determined by Western blotting (**D**). WT and *Aip* KO MEFs were starved of amino acids for 3 hours (EBSS with 1% dialyzed FBS) and stimulated with leucine (0.8mM) and glutamine (4mM) and mTORC1 (pS6K1) activation determined (**E**).

Graphs show the mean ± SEM from at least two independent experiments. Students un-paired, paired *t*-test and 2-way ANOVA were used for analysis

**Supplementary Figure 6**

Relative proportion of amino acids derived from the culture supernatant of WT and *Aip* KO MEFs (**A**). Relative amounts of glycolysis metabolites from the culture supernatant of WT and *Aip* KO MEFs from uniformly labelled (U-^13^C_6_) glucose (**B**). Viability of WT and *Aip* KO MEFs and healthy control (HC) and patient derived dermal fibroblasts cultured in glucose free media over 48 hours (**C**). Viability of WT and *Aip* KO MEFs cultured in the presence of glycolysis inhibitors 2-deoxy-D-glucose (2-DG) (10mM) and oligomycin (Oligo) (1μM) after 48 hours (**D**). Relative amounts of glutamine metabolites from the culture supernatant of WT and *Aip* KO MEFs from uniformly labelled (U-^13^C_6_) glucose (**E**). Graphs show the mean ± SEM from at least two independent experiments. Students un-paired, paired *t*-test used for analysis.

**Supplementary Figure 7**

Relative amounts of TCA cycle intermediates derived from labelled glucose (U-^13^C_6_) (**A-B**) from WT and *Aip* KO MEFs. Relative amount of uniformly labelled glucose (m+3) incorporated into amino acids from WT and *Aip* KO MEFs (**C**). Relative proportion of uniformly labelled glucose in amino acids from WT and *Aip* KO MEFs (**D**). Heatmap of RNA-sequencing expression data of genes involved in the TCA cycle from WT and *Aip* KO MEFs (**E**). Relative expression of electron transport proteins in WT and *Aip* KO MEFs (**F**).

Overview of serine-1 metabolism (**G**). Relative total amounts of serine and glycine and from uniformly labelled (U-^13^C_6_) and the relative incorporation of labelled serine and glycine from WT and *Aip* KO MEFs (**H**). Heatmap of RNA-sequencing expression data of serine-1 metabolism genes in WT and *Aip* KO MEFs (**I**). Results of at least two independent experiments. Graphs show the mean ± SEM from at least two independent experiments. Students un-paired, paired *t*-test, 1-way and 2-way ANOVA were used for analysis.

**Supplementary Figure 8**

Glutamine import and metabolism showing the action of glutaminolysis inhibitors used (**A**). RNA-sequencing expression data of glutaminolysis genes in WT and *Aip* KO MEFs (**B**). Expression of enzymes involved in glutaminolysis from WT and *Aip* KO MEFs (**C**). Viability after 48 hours following treatment of glutaminolysis inhibitors L-DON (1mM), AOA (5mM), BPTES (1μM) and EGCG (50μM) of WT and *Aip* KO MEFs (**D**). WT and *Aip* KO MEFs were treated with uniformly labelled carbon glutamine (U-^13^C_5_) and the relative amounts of TCA intermediates and the relative proportion of oxidative (m+4) and reductive (m+3) glutamine metabolism calculated (**E**). Using uniformly carbon and nitrogen labelled glutamine, the relative proportion of in glutamine metabolites from WT and *Aip* KO MEFs (**F**). Graphs show the mean ± SEM from at least two independent experiments. Students un-paired and paired *t*-test and 1-way ANOVA was used for analysis.

**Supplementary Figure 9**

Comparison of human and zebrafish AIP genomic DNA and AIP structure as determined by Alphafold and Swissfold. The zebrafish AIP protein shares 78.8% identity with the human protein (**A**). Design and generation of a homozygous *aip* mutant zebrafish line. Strategy used to target *aip* in zebrafish using CRISPR/Cas9 (**B**). The 29 bp frame shift caused by the deletion leads to a premature stop codon in the PPIase domain, thereby disrupting the downstream C-terminal tetratricopeptide repeat (TPR) domain. Comparison of yolk size from day 2 to day 6 post fertilization (dpf) in WT and *aip* KO zebrafish (**C**).

**Supplementary Figure 10**

Metabolic analysis of zebrafish. Relative amounts of glycolysis products (**A**), TCA cycle intermediates and amino acids derived from uniformly labelled glucose (U-^13^C_6_) (**B**). Glutaminolysis products derived from labelled glutamine derived from uniformly labelled glutamine (U-^13^C_5_) (**C**). Students un-paired *t*-test and 1-way ANOVA was used for analysis

**Supplementary Figure 11**

Upon birth, neonates need to be able to induce autophagy to adapt to periods of starvation. AIP deficient embryos develop normally, but upon birth have an inability to initiate autophagy and fail to develop. This phenotype was recapitulated in *aip* KO zebrafish that died following depletion of the yolk (**A**). Loss of AIP results in defective proteasome activity, leading to an accumulation of ubiquitylated proteins and a shortage of amino acids (**B**). In AIP deficient cells upon starvation, ATG9a^+^ vesicles that contain the enzyme PI4K3β are retained in the Golgi, resulting in less PI4P and other enzymes and lipids required for autophagosome enlargement and maturation (**C**). AIP deficient cells had defective lysosomes impairing efficient breakdown of proteins to be recycled (**D**). Summary of metabolic reprogramming occurring in *Aip* KO MEFs. Loss of Aip resulted in increased glycolysis, serine-1 metabolism and increased reliance on glutamine metabolism that was used to fuel glycolysis and contribute to serine-1 metabolism (**E**).
